# Integrating intraspecific variation into population dynamics reveals how interacting species persist in mutualistic communities

**DOI:** 10.1101/2024.09.12.612448

**Authors:** Blanca Arroyo-Correa, Ignasi Bartomeus, Pedro Jordano, E. Fernando Cagua, Daniel B. Stouffer

## Abstract

Mutualistic interactions among organisms are fundamental to the origin and maintenance of bio-diversity. Yet, the study of community dynamics often relies on species-level values, ignoring the effects of intraspecific variation. We propose a theoretical framework for evaluating the extent to which variation within populations can influence species’ persistence in mutualistic systems. Next, drawing from detailed empirical data on plant–pollinator interactions and plant fitness, we quantify intraspecific variation in the mutualistic benefits received by plants and incorporate this variation into estimations of structural stability, a robust theoretical measure of species’ likelihood of persistence. Through explicit consideration of intraspecific variation, we demonstrate that, in the absence of heterospecific plant competitors, plant populations composed of specialized individuals promote the persistence of the plant species they belong to and their associated pollinator community. However, these positive effects do not hold when plant species compete with a broader plant community. In this case, having more heterogeneous mixtures of plant individuals (i.e., both specialized and generalized) is more beneficial for a population’s persistence. By integrating the proposed framework with empirical data, we are able to explicitly account for individual-level variation, opening the door to a better understanding of the mechanisms promoting biodiversity in mutualistic communities.

## INTRODUCTION

Ecological interactions, such as those involving pollination, seed dispersal or herbivory, have been shown to play a key role in shaping community assembly and ecosystem functioning (Bascompte & Jordano, 2007; Elton, 1927). Even though these interactions are usually summarized at the species level (Vázquez *et al*., 2009), natural populations often display intraspecific differences in traits among individuals and thus exhibit differences in the interactions those individuals can potentially establish (Siefert *et al*., 2015; Van Valen, 1965; Van Valen & Grant, 1970). Most populations are in fact composed of heterogeneous collections of specialized individuals, which interact only with a small portion of the available partners, and generalized individuals, which interact with a larger portion of these partners (Araújo *et al*., 2008; Arroyo-Correa *et al*., 2023; Bolnick *et al*., 2003; Violle *et al*., 2012).

Despite the fact that intraspecific variation is both well documented (Bolnick *et al*., 2003; Violle *et al*., 2012) and expected to influence species coexistence (Hart *et al*., 2016; Stump *et al*., 2022), its effects on species persistence remain unclear. This is likely due to both the difficulty of obtaining the necessary fine-scale data, but also because of the current limitations of theoretical approaches. In competitive contexts, several conceptual works have suggested different mechanisms through which intraspecific variation can influence community dynamics (Barabás & D’Andrea, 2016; Bolnick *et al*., 2011; Hart *et al*., 2016). For example, within species differences in competitive ability can ‘blur’ between species differences in competitive ability, and therefore individual variation is expected to break down competitive hierarchies, promoting coexistence (Clark *et al*., 2010; Fridley *et al*., 2007; Hubbell, 2005). In contrast, classical niche theory posits that intraspecific niche variation might lead to an increase in species-level niche overlap, diminishing the probability of species coexistence (Doebeli, 1996; Roughgarden, 1972; Slatkin, 1980).

Unlike competitive communities, we know substantially less about how intraspecific variation affects the persistence of mutualistic assemblages, despite their standing as the cornerstone of many ecological systems (Bastolla *et al*., 2009; García *et al*., 2018; Winfree, 2013). In mutualistic plant–animal communities, combinations of specialized and generalized individuals (in terms of resource use, e.g., pollinators used by plants, fruits used by seed dispersers) are expected to strongly impact dynamics through effects on fitness and niche differences between species. For instance, in plant–pollinator assemblages, if different individuals of the same plant species vary in their attractiveness to different pollinators, these plant individuals may also differ in the fitness benefits received from the pollinators with which they interact (Arroyo-Correa *et al*., 2021). Generalized plant individuals are likely to interact with more beneficial pollinators merely by attracting a wider range of pollinator species, leading to more successful fitness outcomes on the whole (Arroyo-Correa *et al*., 2021; Gómez *et al*., 2007). Therefore, the extent to which co-occurring plant species are composed of more generalized or more specialized individuals is likely to determine differences between co-occurring plant species in fitness. Plant individuals’ specialization/generalization in pollinator use can also shape the patterns of pollinator sharing among plant species, thereby creating indirect plant–plant interactions, which are also predicted to affect the scope for coexistence within communities (Bastolla *et al*., 2009). Through these effects, we expect intraspecific differences in plant generalization to impact the persistence of mutualistic systems and hence, the ability of species to sustain themselves within the community. Therefore, acknowledging a connection between intraspecific variation in mutualistic assemblages, either within one guild (e.g., plants) or across multiple guilds (e.g., plants and pollinators), and community dynamics may advance our understanding of the mechanisms fostering diversity.

A necessary condition for all species to persist in a community is that those species can maintain positive abundances. Using population-dynamics models, we can quantify the range of intrinsic growth rates that can fulfill that condition (Grilli *et al*., 2017; Saavedra *et al*., 2016b; Song *et al*., 2018). This range is called the feasibility domain, and can be thought of as the set of environmental conditions under which all co-occurring and interacting species in a given site and time can exhibit positive abundances (Cervantes-Loreto *et al*., 2023; Rohr *et al*., 2014; Song & Saavedra, 2018b). In a nutshell, this “structural stability approach” posits that the larger the feasibility domain, the greater the opportunities to coexist under random variation in intrinsic growth rates. Despite its proven power to predict species persistence (Domínguez-García *et al*., 2024), this approach has yet to be employed to explore how observable variation among individuals may influence the dynamics of ecological assemblages.

Here, we demonstrate how to expand the structural stability framework to incorporate intraspecific (i.e., within-population) variation in ecological interactions to the population dynamics of plant– animal mutualistic communities. Our proposed framework serves as a tool for evaluating the extent to which various forms of within-population variation could influence the emergent dynamics of these communities, and the range of conditions under which species can be sustained. After presenting the underlying theory, we work through an empirical case study to illustrate how observed intraspecific plant variation in pollinator use influences species’ persistence in plant–pollinator communities. First, we empirically estimate the mutualistic benefits received by plant individuals from different pollinator species, and used this information to parameterize plant intraspecific variation in the proposed theoretical framework. Second, we assess how intraspecific variation in pollinator use can impact the feasibility of mutualistic assemblages. Inspired by the results of this analysis and also by a theoretical testing of the proposed framework, we next evaluate how the proportion of specialized individuals (i.e., those interacting with few pollinator species) within a single plant species influences the feasibility of its mutualistic assemblage in the absence of heterospecific plant competitors. Further, we also investigate whether a high level of individual specialization in focal plant species can influence their ability to persist when they are embedded in the broader plant community.

## THEORETICAL FRAMEWORK

### Population dynamics in mutualistic communities

The dynamics of plant and animal populations in a community composed of multiple plant species that interact mutualistically with multiple animal species can be approximated using a generalized Lotka–Volterra system of the form:

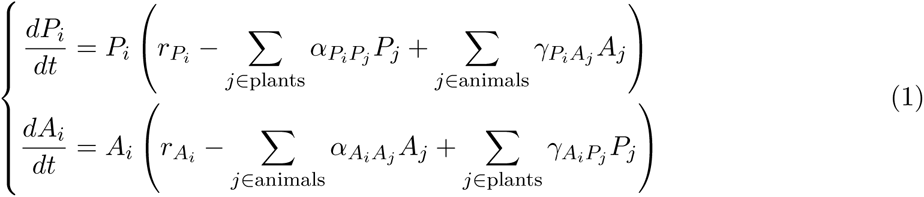

where *P_i_* and *A_i_* denote the abundance of plant species *i* and animal species *i*, respectively. The parameters of this mutualistic model correspond to the populations’ intrinsic per capita growth rates (*rP_i_* and *rA_i_*, which represent how plant species *P_i_* and animal species *A_i_* grow in isolation), within-guild interactions (*αP_i_P_j_* and *αA_i_A_j_*), and between-guild interactions (*ϒP_i_A_j_*, *ϒA_i_P_j_*). These within-guild or between-guild interactions represent per-capita biotic effects of species *P_j_* or *A_j_* on the per-capita growth rates of species *P_i_* or *A_i_*. Formally, plant and animal species’ intrinsic growth rates are not explicitly constrained by sign restrictions biologically and in the formulation. Meanwhile, *α* coefficients must be positive in the formulation, reflecting the negative effects of within-guild competition on population growth; and *ϒ* coefficients must also be positive, in order to reflect the fact that between-guild mutualistic interactions lead to a positive effect on the growth rate of the interacting species. Hereafter, we will use the term ‘species’ to refer to all individuals that belong to the same single-species population. In a community comprising two plant species and two animal species, the collective population dynamics can be expressed in matrix form as follows:

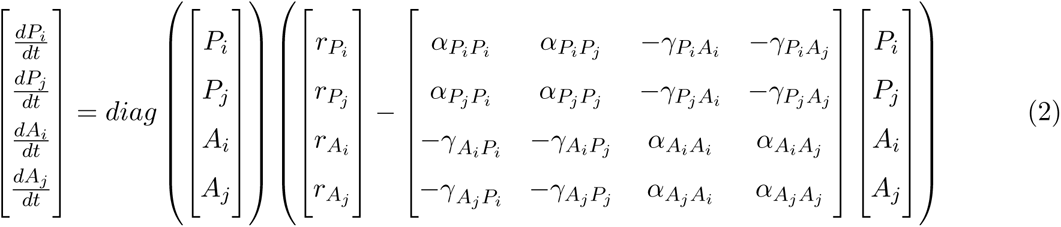

where *diag*(**x**) creates a diagonal matrix with values from the vector **x** on the diagonal and zeros everywhere else.

### Incorporating within-population variation

Any given species might be potentially composed of individuals or groups of individuals that differ in ecologically meaningful ways. Here we assume that individuals act as representative subsets of the larger population (‘individual types’ hereafter), similar to ‘phenotypes’ in eco-evolutionary contexts (Maynard *et al*., 2019). For the sake of simplicity, we will start by considering how to specify the population dynamics of a community solely consisting of two different individual types, *s* and *t*, within a single plant species *i* which interact with two different individual types, *s* and *t*, within a single animal species *i*. The population dynamics of such a community can thus be described as:

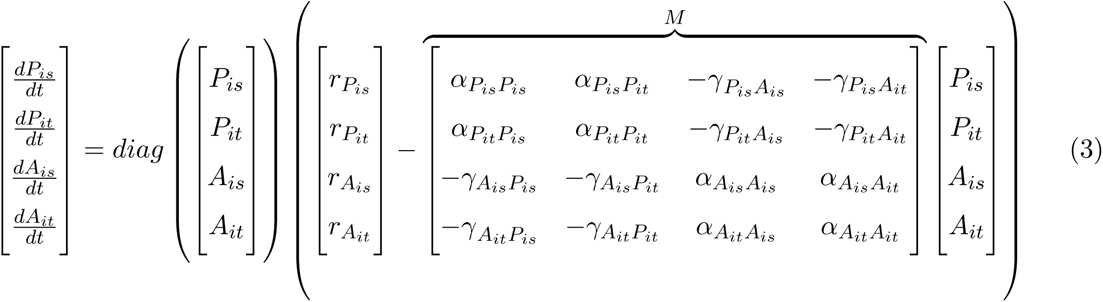

where within-population variation is enabled by the fact that within-type growth rates (e.g., *rP_is_* and *rP_it_*) and the within-type, between-type, and across-type interaction coefficients may all take distinct values (e.g., *ϒP_is_A_is_* may differ from *ϒP_it_A_is_*, and so on).

### Feasible conditions

If we call the vector of individual types’ growth rates 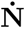, the vector of intrinsic growth rates **r**, the vector of total individual types’ abundances **N**, and the type-by-type interaction matrix *M*, we can more compactly write the population-dynamics equation as:

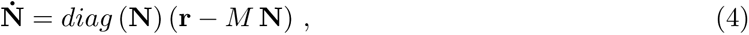

and the equilibrium condition is given by **r** = *M* **N***^*^* .

Given only information about the interaction strengths in *M*, the “unknowns” in the equilibrium condition for this model are **r** and **N***^*^*. The core idea underpinning the feasibility domain is to explore whether *M* imposes strong or weak constraints on species’ abilities to sustain positive equilibrium abundances (Rohr *et al*., 2014; Saavedra *et al*., 2013; Song *et al*., 2018). By restricting ourselves to any and all feasible equilibria (i.e., vectors **N***^*^* containing only positive values), we shift **N***^*^* into the “knowns” category. Consequently, one can mathematically define the feasibility domain as the set of vectors **r** for which all equilibrium abundances in **N***^*^* = *M^−^*^1^ **r** are positive, where *M^−^*^1^ is the inverse of the matrix *M* . Here, it is also possible to search for growth rate vectors that, in addition to ensuring feasibility, are subject to further biotic constraints based on the biological characteristics of the study system (see below for the implementation of biotic constraints in our specific case study).

## EMPIRICAL CASE STUDY

The theoretical framework proposed above explicitly incorporates variation within populations when describing the dynamics of mutualistic communities. To illustrate how empirically observable intraspecific variation in ecological interactions modulates the dynamics of a real mutualistic community, we used finely resolved data of pollinator visitation to plant individuals and plant fitness proxies measured on these same individuals. First, we empirically estimated the mutualistic benefits given by different pollinator species to each plant individual by integrating multiple sources of intraspecific variation across plants (pollinator assemblage, visitation rates and flower production) and fitness data (fruit and seed production). Second, we used these estimates to characterize variation among plant individuals in the mutualistic benefits they received, and therefore, in how they are expected to contribute to the dynamics of their plant populations. With the parameterized model, we explored how this intraspecific variation within each focal plant species can influence the size of the feasibility domain of their populations’ mutualistic assemblages in the absence of heterospecific plant competitors (i.e., the ability of these species to sustain themselves as well as their pollinators). Inspired by the results of this analysis and also by a theoretical testing of the proposed framework, we then evaluated the effects of plant individual specialization on pollinator use in the absence of heterospecific plant competitors, and further, when they are embedded in the broader plant community.

### Study system

Here we focus on three representative plant species sampled in Mediterranean shrublands of Doñana National Park, located on the Atlantic coast of southwestern Spain (37°00’29.736"-N –6°30’24.919"-W, 25 m a.s.l.). These focal species belong to the Cistaceae family and flower consecutively throughout the flowering period of the plant community (starting with *Halimium calycinum*, followed by *Cistus libanotis* and ending with *Halimium halimifolium*), overlapping with each other to varying extents. Plant individuals of these focal species were sampled in six different study plots within an area of one square kilometer to capture environmental heterogeneity. We recorded pollinator visits to these plant individuals during the 2021 peak flowering season, between early February and mid July. We recorded pollinator visitations to 73, 86 and 84 plant individuals belonging to *Cistus libanotis*, *Halimium calycinum* and *Halimium halimifolium*, respectively. These plant individuals were selected following a stratified random sampling scheme within each study plot to capture within-plot heterogeneity. Pollinators were considered as all insects touching flowers’ reproductive structures, and were identified as morphospecies (‘species’ hereafter). We observed a total of 55 pollinator species visiting *Cistus libanotis*, 33 visiting *Halimium calycinum*, and 38 visiting *Halimium halimifolium*. On average, interaction sampling completeness was above 85% for all plant individuals sampled (Arroyo-Correa *et al*., 2023). For each focal plant individual, we used these records to estimate the frequency of interactions per minute established by each pollinator species along the entire flowering season (Arroyo-Correa *et al*., 2023), and we also measured two proxies of female plant fitness at the flower level: fruit set (i.e., proportion of flowers setting final-sized fruits) and seed set (i.e., number of seeds per flower). All plant species produced multiple inflorescences and multiple flowers per inflorescence. We randomly selected five inflorescences per plant individual and estimated whether flowers produced at the beginning of the season in these inflorescences resulted in any fruit production at the end of the season. The total number of sampled flowers was 3017, 6229, and 7127 for *Cistus libanotis*, *Halimium calycinum*, and *Halimium halimifolium*, respectively. To estimate seed production in each plant, we randomly sampled a maximum of 10 fully developed fruits from different randomly selected inflorescences to obtain the number of seeds per fruit. We obtained the seed count from 730, 820 and 732 fruits of *Cistus libanotis*, *Halimium calycinum*, and *Halimium halimifolium*, respectively. Each week along the flowering season, we also counted the number of individual flowers in each plant individual and summed the weekly number of flowers as an estimate of the total number of flowers produced by a plant individual.

All these plant species were composed of a mixture of plant individuals ranging from more specialized (i.e., interacting with a small proportion of the available pollinator species) to more generalized (i.e., interacting with a large proportion of the available pollinator species) (Arroyo-Correa *et al*., 2023). We considered that the plant individuals sampled in the field are representative types of individuals within their populations (i.e., proper subsets spanning the whole range of individual variation), and therefore we used each of these individuals to characterize a separate ‘individual type’ in our theoretical framework.

### Theoretical framework adapted to the empirical case

We unified our empirical data with the proposed theoretical framework to describe the population dynamics of plant species composed of different plant individuals, which act as representative individual types, that differ in the interactions they establish with pollinator species. Given that we only had data on intraspecific variation on the plant side, we adopted a “mixed” perspective with equations defined at the individual type level on the plant side and at the species level on the pollinator side. When exploring the feasibility domain in this specific case, we search for intrinsic growth rate vectors that, in addition to feasibility, are subject to two further specific biotic constraints: one on the pollinator side and one on the plant side.

On the pollinator side, we restrict ourselves to only feasible equilibria that are consistent with pollinator species’ observed relative abundances. We used the relative visitation rates of pollinator species as a proxy for their relative abundances, assuming that pollinators’ visitation rates are proportional to abundance. We primarily include this constraint on the pollinator side for biological reasons. First, in doing so, we are *conditioning* subsequent calculations of the feasibility domain on the actual pollinator community that the plant populations had available to them in the field. Therefore, this allows us to focus solely on the growth rate vectors (**r**) whose corresponding feasible equilibria are consistent with this additional biotic constraint (Cervantes-Loreto *et al*., 2023; Song *et al*., 2018). Second, by imposing this constraint we can also better isolate the plant-specific impacts of intraspecific variation on the size of the feasibility domain (our primary research question). As a mathematical benefit, to make this constraint work we only require a single unknown intrinsic growth rate on the pollinator side to be paired with a single “known” total pollinator abundance which simplifies the estimation of the size of the feasibility domain.

On the plant side, we restrict ourselves to growth rate vectors for which individual types belonging to the same species are constrained to exhibit identical intrinsic growth rates (i.e., *rP_is_* = *rP_it_* when *s* ≠ *t*). In first instance, we impose this constraint because we are primarily interested in understanding the implications of different individual types engaging in different interactions and consequently making different net contributions to the maintenance of their populations as a whole. Moreover, empirical evidence suggests that variation in intrinsic growth rates is not a good predictor of individual fitness and that variation in interaction coefficients often has a greater influence on community dynamics than variation in intrinsic growth rates (Bartomeus *et al*., 2021; Edmunds, 2017; Gibert *et al*., 2016; Hart *et al*., 2019; Levine & HilleRisLambers, 2009; Nguyen *et al*., 2024; Sibly & Hone, 2002). Lastly, we impose this constraint because, without it, we might otherwise end up with a feasibility domain composed of growth-rate vectors with greater variation within species than between species, which we expect is not biologically realistic. Indeed, without imposing this constraint there is nothing mathematical that would otherwise distinguish between two individual types that belong to the same species *versus* two individual types that belong to different species.

To achieve all of this mathematically, we adapted the framework to our empirical case combining relevant elements of Eqs. 3 and 2. For example, to consider a community composed of a single plant species whose population comprises two different types of individuals and its two interacting pollinator species, the dynamics of this community can be described as:

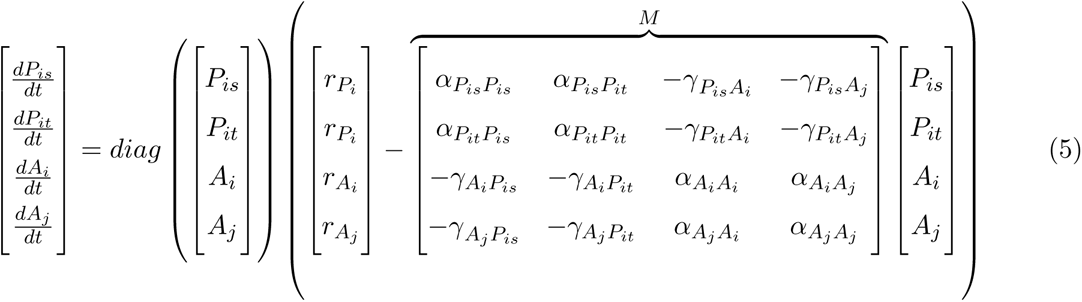

Here, intraspecific plant variation (i.e., within-population variation) is represented as variation in coefficients *ϒP_is_A_i_* across different types of plant individuals belonging to the same species. Note that, though individual types *P_is_* and *P_it_* share the common growth rate parameter *rP_i_*, this in no way implies that they will have the same equilibrium density. Indeed, variation in the mutualistic benefits they receive from their different interacting pollinators (i.e., *ϒP_is_A_i_* vs *ϒP_it_A_i_*) will create variation in their equilibrium relative abundances, and thus also in how each type contributes to the maintenance of the population of *P_i_*as a whole.

Importantly, after imposing our constraints (see Appendix A), feasibility of the vector **N***^*^* is still measured in terms of growth rates that vary at the species level (i.e., at the level of the overarching populations *P_i_*, *P_j_*, *A_i_*, *A_j_*, etc.) even though interactions vary within, across, and between individual types.

In the most general case, the domain of intrinsic growth rate vectors (**r**) leading to positive species abundances (**N***^*^*) is geometrically described by an algebraic cone, where the borders are established by the column vectors of the interaction matrix *M* (Logofet, 1993; Ribando, 2006). The normalized solid angle of this cone can be interpreted as the probability that a randomly sampled a vector of intrinsic growth rates falls inside that cone. With our additional relative abundance and growth rate constraints, the space of feasible growth rate vectors is no longer guaranteed to be conic, and analytical approaches to estimate the magnitude of the solid angle no longer hold (Song *et al*., 2018). We therefore adopt a fully Monte Carlo approach (Song & Saavedra, 2018a) to estimate the proportion of randomly sampled vectors of intrinsic growth rates that are consistent across all of the aforementioned constraints. In all cases, the higher this probability for a given community, the higher its potential to accommodate variations in species growth rates while maintaining feasibility.

### Parameterization of interaction coefficients

We focused on evaluating how the empirically observed intraspecific variation in the mutualistic benefits received by plant individual types influences the ability of species to persist. To this end, we empirically parameterized the mutualistic benefits provided by each pollinator species to each plant individual type, denoted in the model as *ϒP_is_A_i_* . This parameterization was based on our empirical data on plant fitness proxies (i.e., proportion of flowers setting fruit and number of seeds produced per fruit), visitation rates and flower production (Box 1).

To maintain our focus on the effects of the observed intraspecific plant variation in the mutualistic benefits received, all other coefficients in the interaction matrix *M* (*αP_is_P_is_*, *αP_is_P_it_*, *αA_i_A_i_*, *αA_i_A_j_ ϒA_i_P_is_*, and *ϒA_i_P_it_*) were parameterized following a mean-field approximation (Appendix D), as already done in previous works (García-Callejas *et al*., 2023; Rohr *et al*., 2014; Song *et al*., 2018).

### Within-population plant variation in mutualistic benefits

To visualize how different plant fitness components (Box 1) ultimately combine together to determine the number of seeds that a given plant individual contributes to the plant population’s seed bank, we performed a principal component analysis (PCA) for each of our three focal plant species. In addition to the three components influencing plant fitness that are included in Eq. 8, the total mutualistic benefits received by plant individuals may also be influenced by the degree of these plant individuals (i.e., the richness of pollinator species with which they interact). The degree of plant individuals may increase the overall contribution of pollinators to a given plant individual’s fitness (i.e., the sum of the contributions *βP_is_A_i_* of all pollinator species to a given plant). This procedure resulted in an ordination of plant individuals in relation to the multivariate space defined by these multiple components influencing plant individuals’ fitness. This ordination allowed us to evaluate how the combination of these components modulates fitness variation among plant individuals within a population, and therefore intraspecific variation in the mutualistic benefits received by plant individuals.

### Impact of within-population plant variation on feasibility

To explore the implications of the presence of intraspecific variation in our empirical data, we examined a scenario where each “individual type” sampled in the field was treated as the sole representative of its population. This scenario represents the hypothetical situation, for example, where we only had empirically observed interaction data for a single plant individual. This approach allowed us to isolate the effect of intraspecific variation by creating populations composed of a single type of individual. For each of these mutualistic assemblages in which the plant population only comprised a single plant individual type, we estimated the size of the feasibility domain. In addition, we assessed whether those feasibility estimates were correlated with attributes of those plant individual types (i.e., fitness components). This, in turn, provided a baseline for assessing the influence of this variation in our empirical data on the size of the feasibility domain when plant populations are composed of mixtures of individual types.

### Consequences of individual specialization within a single plant population

We aimed to evaluate how the inferred intraspecific plant variation in mutualistic benefits received (*ϒP_is_A_i_*) affected the feasibility of a system composed of multiple individual types from a single plant species and their interacting pollinator community, in the absence of heterospecific competitors. Before performing such an evaluation directly using our empirical data, we first applied the adapted mathematical framework (Eq. 5) to a simplified system, aiming to establish theoretical expectations for the effects of intraspecific plant variation on the collective feasibility (Box 2). This theoretical exploration revealed that the level of specialization strongly influences the size of the feasibility domain in systems where plant populations are composed of multiple individual types. Building on this insight, we assessed how a plant population dominated by specialized individual types affects the feasibility of the collective system (i.e., the plant population together with its pollinator assemblage) compared to a plant population with mixtures of specialized and generalized plant individual types. This contrast was also motivated by the fact that pressures imposed by global change are likely to exacerbate the risk of extinction for large individuals, especially for plants (Bennett *et al*., 2015; Gora & Esquivel-Muelbert, 2021), which often coincide with those exhibiting generalized resource use patterns. Thus, size-structured induced mortality might lead to the emergence of populations composed of smaller individuals, restricted to highly specific local environments or exhibiting more specialized traits, which are usually more specialized in resource use. Consequently, we considered that an increased individual specialization level within populations was the most plausible future scenario under global change. To address our main questions, we rely on the definition of specialized and generalized plant individuals based on their richness of pollinator visitors (i.e., degree), such that individuals range between more specialized (lower degree) to more generalized (higher degree) on pollinator use.

For each plant species separately, we created interaction matrices composed of multiple plant individual types and the pollinator species visiting the plant species to which these individuals belong, over an increasing number of plant individual types (i.e., hereafter “plant population size”). In a first scenario, plant population size was increased by progressively including plant individuals in order of increasing degree (i.e., from the most specialized to the most generalized individual types). To break ties, plant individual types with the exact same degree were incorporated based on decreasing specificity (i.e., decreasing coefficient of variation of interactions using visitation rates, Poisot *et al*., 2012). In a second scenario, we increased plant population size by progressively including plant individuals selected at random (N=100 iterations), regardless of their degree. In each scenario, and for each interaction matrix composed of the plant population (differing in the types and the number of types included) and its pollinator species, we estimated the size of the feasibility domain. Furthermore, for the feasibility domain estimated for each interaction matrix, we also partitioned the effects on the plant and pollinator sides by estimating the range of feasible growth rates for the plant population and pollinator species, respectively. Therefore, for a given plant population size, differences in the feasibility domain size and in the ranges of feasible growth rates for plants and pollinators between the first and second scenarios represent the consequence of the focal plant population being composed a higher proportion of specialized individuals compared to a more heterogeneous mixture of individuals.

### Effects of within-population individual specialization across the plant community

After exploring how individual specialization in a single plant species affects the persistence capacity of its mutualistic assemblage, we aimed to assess the effects of individual specialization within a focal plant species in the presence of heterospecific competitors (i.e., when it is embedded in the broader plant community). We estimated these effects as changes in the ‘vulnerability’ of plant species within the community (Allen-Perkins *et al*., 2023; Lepori *et al*., 2024; Medeiros *et al*., 2021). To estimate vulnerability, we first sampled vectors of growth rates in the feasibility domain, subject to our specified constraints. Each of these vectors corresponds to a vector of feasible individual type densities. Second, we obtained species feasible density vectors by summing feasible densities across plant individual types belonging to the same species. Then, we calculated the normalized density for each plant species and each vector by dividing its density within a given vector by the maximum density of that species observed across all vectors. The vulnerability of a given species is calculated as the proportion of feasible vectors of species densities where that species is the one closest to the boundary of exclusion (i.e., where it exhibits the lowest normalized density). This vulnerability metric ranges between 0 and 1, and quantifies the average probability of that a given species would be the first one excluded if a random perturbation in intrinsic growth rates took place in a feasible community under generalized Lotka–Volterra dynamics.

We used the same scenarios as mentioned above for the plant population-level analyses but focused on focal populations comprising a fixed number of individual types. In the first scenario, the plant community was composed of the N=40 most specialized plant individuals of the focal plant species and all plant individuals belonging to the other two species. In the second scenario, the plant community was composed of N=40 randomly selected plant individuals (100 replicates) of the focal plant species and all plant individuals belonging to the other two plant species. We calculated the ‘vulnerability’ (v) of all three plant species in both scenarios. The difference in this metric between both scenarios, represented as Δv, indicates the effects of a higher proportion of specialized individuals within the focal species on itself and any other plant species in the community. Positive values of Δv indicate that having more specialized individuals of a focal species increases the vulnerability of a given species. Meanwhile, negative values of Δv indicate that having more specialized individuals of a focal species decreases the vulnerability of a given species.

All analyses were performed in R v4.3.1 (R Core Team, 2023).

## RESULTS

### Within-population plant variation in mutualistic benefits

Empirically observed differences among plant individuals in pollinator use translated into variation in plant fitness merely due to differences among pollinator species in their fitness contribution to plants (Fig. 2A, Appendix F). Moreover, the level of this intraspecific fitness variation driven purely by differences in plant individuals’ pollinator assemblages changed when also incorporating intraspecific variation in flower production and visitation rates (Fig. 2B-C). A principal component analysis (PCA) for each of our three focal plant species revealed that these different sources of intraspecific variation explained most of the variance in realized plant individual fitness, expressed as the total number of seeds produced per plant individual (Fig. 2D). The first principal component (PC1) explained 48.11 ± 5.94% [mean ± SD across plant species] of intraspecific variation in fitness resulting from mutualistic interactions. Meanwhile, the second principal component (PC2) explained 26.79 ± 3.44% across plant species.

**Figure 1:**
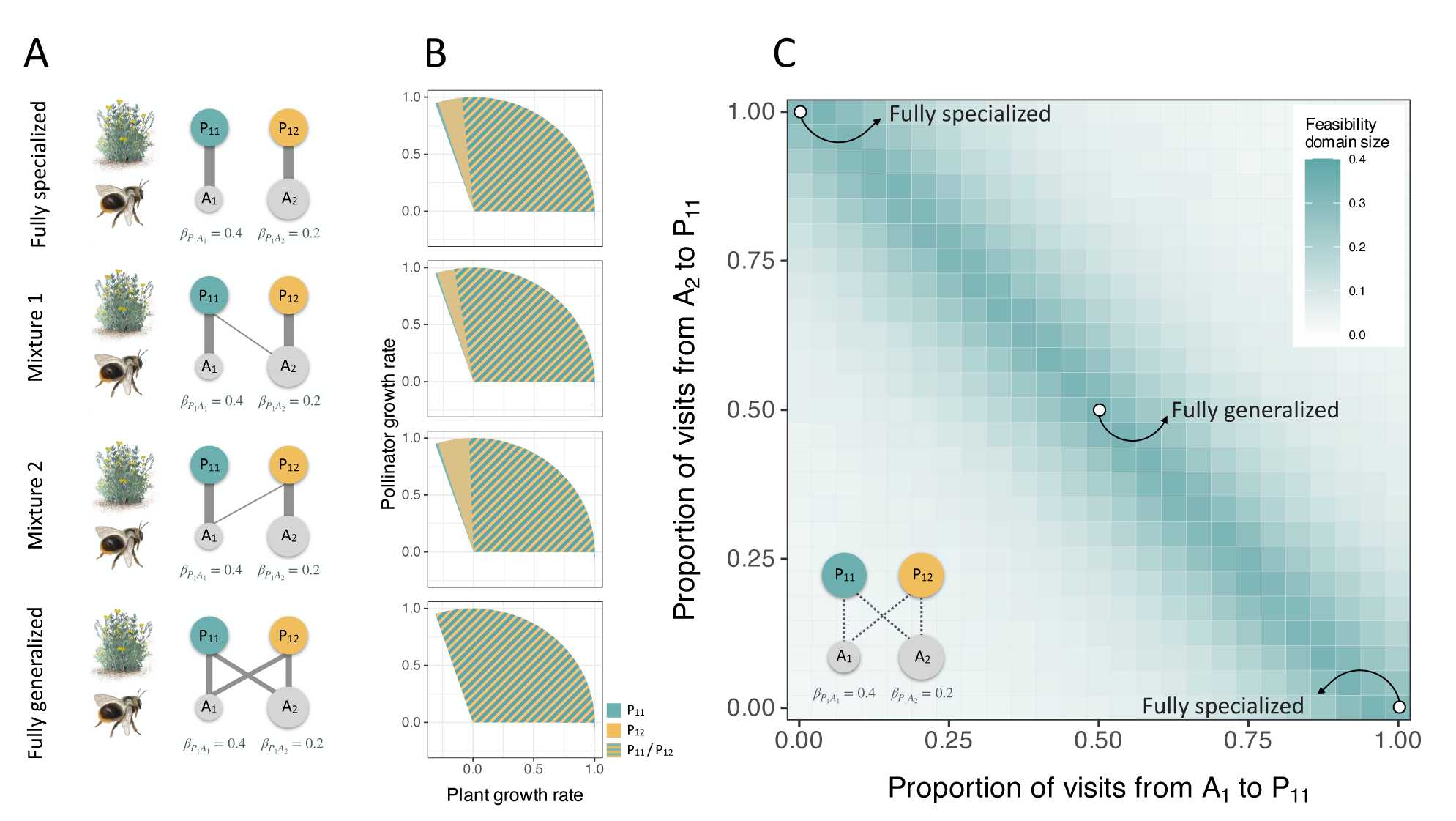
Conceptual overview of how intraspecific plant variation can impact species persistence in a plant–pollinator assemblage. (A) Illustration of a simplified community composed of two individual types belonging to the same plant species (*P*_11_ and *P*_12_, colored nodes), which interact (gray links) with two pollinator species (*A*_1_ and *A*_2_, light gray nodes) that differ in their contribution to plant fitness (i.e., different *β* values) and their relative abundance (indicated by pollinator node size). We considered four ways that the plant species could exhibit different extents of intraspecific variation: ‘fully specialized’, where both plant types are wholly specialized on different pollinator species, ‘mixture 1’ and ‘mixture 2’, where only one of the two types is generalized, and ‘fully generalized’, where both types are generalized. To visually emphasize the differences across cases, link width is shown proportional to visitation rates. (B) For each case, the striped area represents the size of the feasibility domain for the population composed of both plant types, and the blue and yellow areas represent the size of the feasibility domain when the population is only composed of individuals belonging to one of types *P*_11_ or *P*_12_, respectively. Note that in the ‘fully generalized’ case all three areas are equivalent as there is no intraspecific variation. (C) Variation in the feasibility domain size depends on the proportion of visits from both pollinator species (*A*_1_ and *A*_2_) to plant individual type *P*_11_ (dotted links in the inset). The ‘fully specialized’ case is represented in the top left corner, where *P*_11_ is specialized on *A*_2_ and *P*_12_ on *A*_1_, or the bottom right corner, where *P*_11_ is specialized on *A*_1_ and *P*_12_ on *A*_2_. The ‘fully generalized’ case is represented in the center of the grid, where both plant types receive the same exact proportion of visits. The ‘mixture 1’ case is represented by a range of cells where *P*_11_ is generalized and *P*_12_ is specialized on *A*_2_ (spanning varying proportions of visits). The ‘mixture 2’ case is represented by a range of cells where *P*_11_ is specialized on *A*_1_ and *P*_12_ is generalized (spanning varying proportions of visits). Note that all other cells in the grid represent intermediate cases between mixtures of generalized and specialized types and the ‘fully generalized’ case.

**Figure 2:**
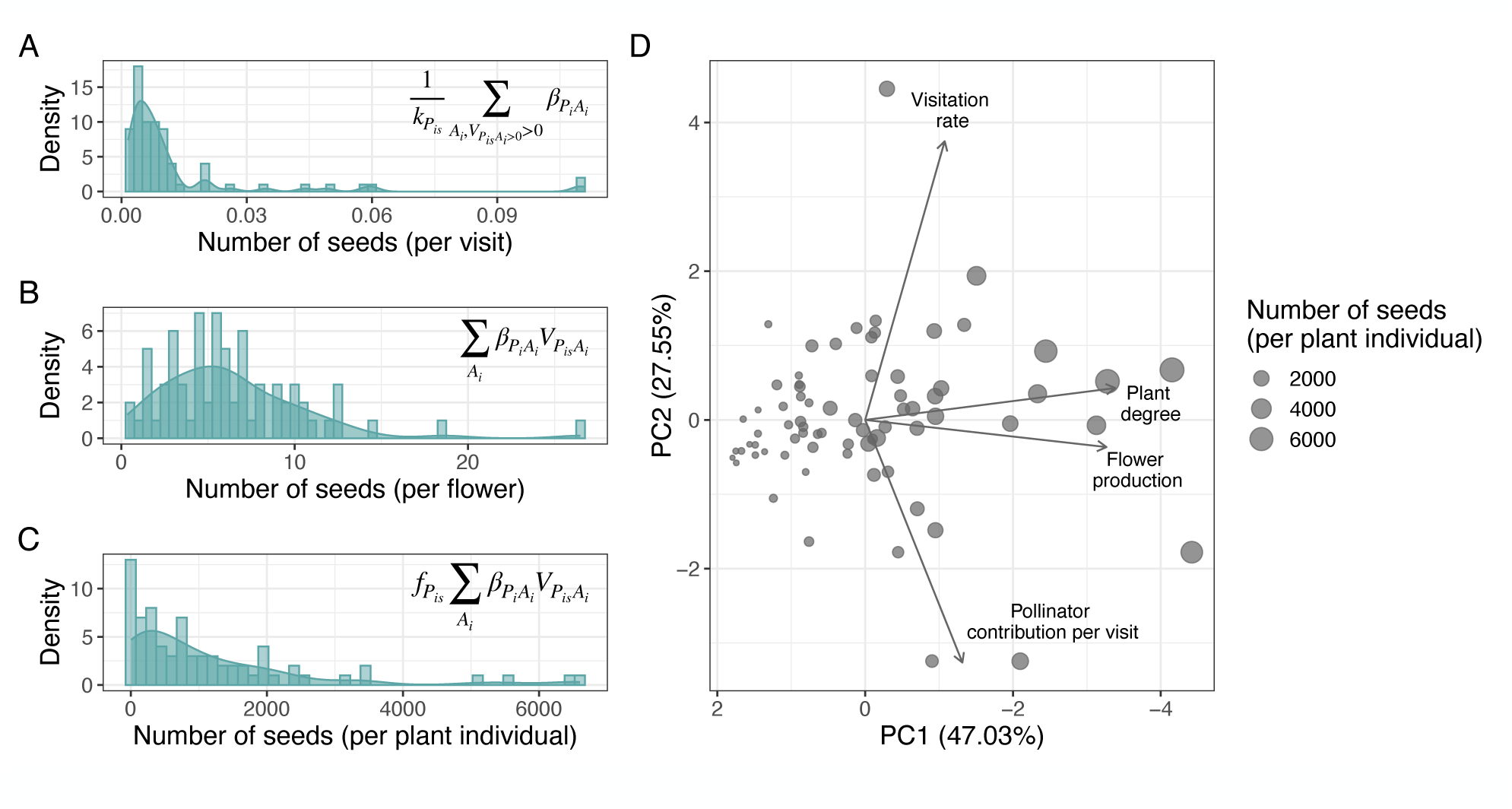
Sources of intraspecific variation influencing plant fitness. Intraspecific plant fitness variation within one of the study species (*Cistus libanotis*) when incorporating different components influencing plant fitness. (A) Histogram of the average number of seeds produced per visit across plant individuals, where intraspecific variation is promoted by differences in pollinator assemblages and their associated contribution to fitness. (B) Histogram of the number of seeds produced per flower across plant individuals, where intraspecific variation is promoted by differences in visitation rates in addition to differences in pollinator assemblages. (C) Histogram of the total number of seeds contributed to the population seed bank across plant individuals, where intraspecific variation is produced by differences in pollinator assemblages, visitation rates and flower production. (D) Principal Component Analysis (PCA) of plant individuals (dots) based on different sources of intraspecific variation influencing plant fitness. Dot size is proportional to the number of seeds produced and contributed to the seed bank of the population. Estimates of the number of seeds per visit (A), per flower (B) and per individual plant (C) were calculated following the equations shown in each panel, in which *kP_is_* denotes the degree of plant individual *P_is_*, *βP_is_A _i_* denotes the number of seeds per flower produced in plant species *P_i_* per visit of pollinator species *A_i_*, *vP_is_A_i_* is the visitation rate per flower of pollinator species *A_i_* to plant individual *P_is_*, and *fP_is_* represents the number of flowers produced by plant individual *P_is_*. We found similar patterns for other focal plant species (Appendix F).

### Impact of within-population plant variation on feasibility

We found that intraspecific variation can significantly alter estimates of the feasibility domain size, even in populations composed of only one individual type (Fig. 3). This result highlights how the feasibility of a system strongly depends on the representative individual type composing the plant population when there is not intraspecific variation. Feasibility estimates consistently varied with individual-level degree, emphasizing the role played by individual-level specialization—defined as the number of pollinator species interacting with each plant individual type—in permitting species persistence. We also found that these feasibility estimates when populations were composed of a single individual type were also associated with other attributes of these individual types (Figs. S10-S12), such as the number of flowers produced. Nevertheless, these attributes were themselves also highly correlated with degree (Figs. 3, S8, S9).

**Figure 3:**
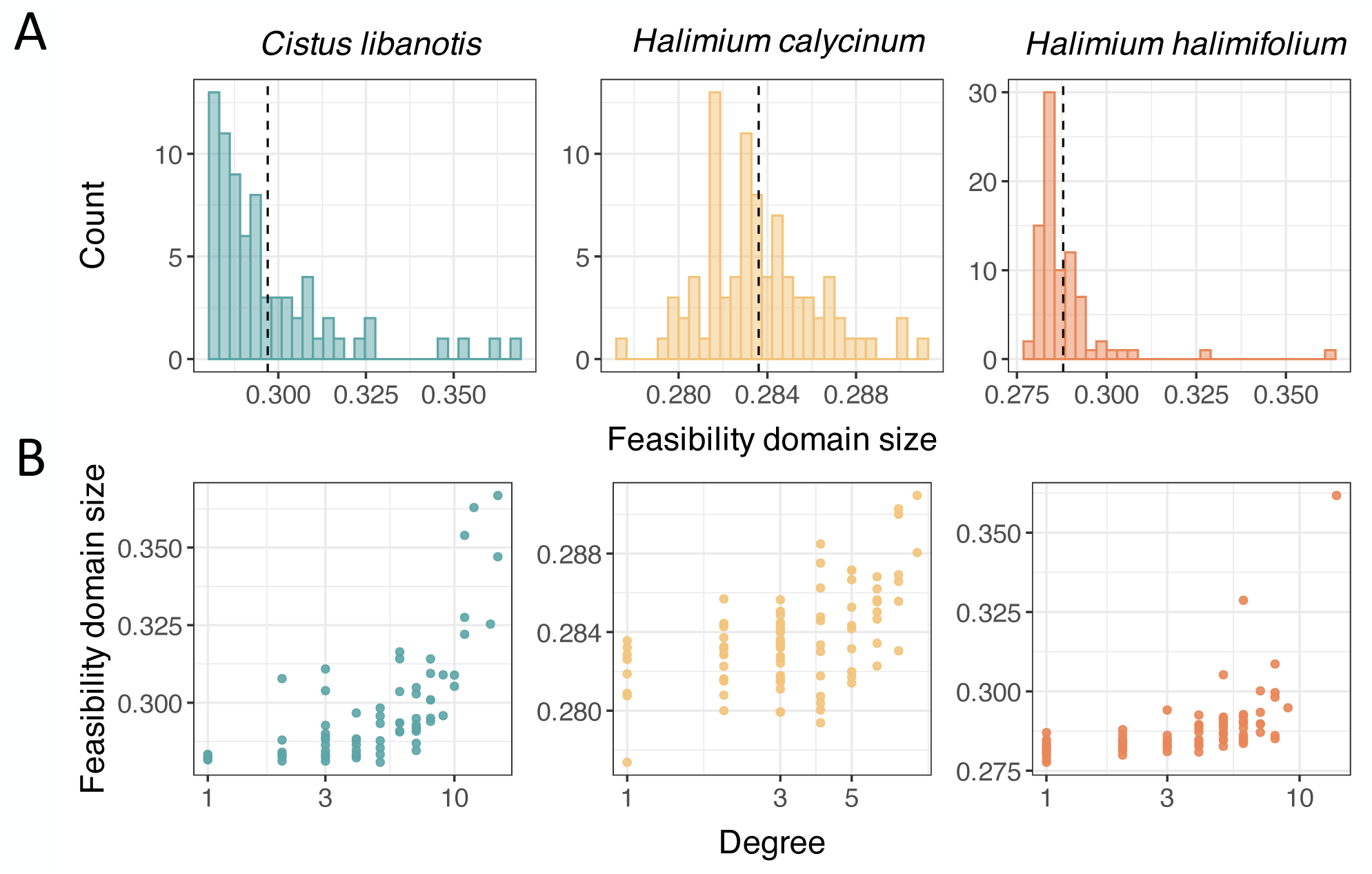
(A) Histograms of the size of the feasibility domain of systems consisting of a plant population and its associated pollinator species, when a single plant individual type is representative of the entire population (i.e., no intraspecific variation). The vertical dotted line represents the averaged feasibility domain across all populations composed of a single individual type. (B) Relationship between the feasibility domain size of these systems and the degree of the individual type (i.e., the number of pollinator species it interacts with) representative of the plant population. Different colors represent different focal plant species. Note the log scale in the x-axis.

### Consequences of individual specialization within a single plant population

Considering each plant population separately and in the absence of heterospecific plant competitors, we found that having a higher proportion of specialized plant individuals within a population increased the feasibility of the population’s mutualistic assemblage, compared to a scenario composed of a mixture of specialized and generalized individuals (Fig. 4A). This positive effect of specialized plant individuals on the feasibility of the entire system was driven by an increase in the range of feasible growth rates on both the plant and pollinator sides (Fig. 4B-C). However, the range of plant growth rates declined more rapidly than that of pollinators as plant population size increased (Fig. 4C).

**Figure 4:**
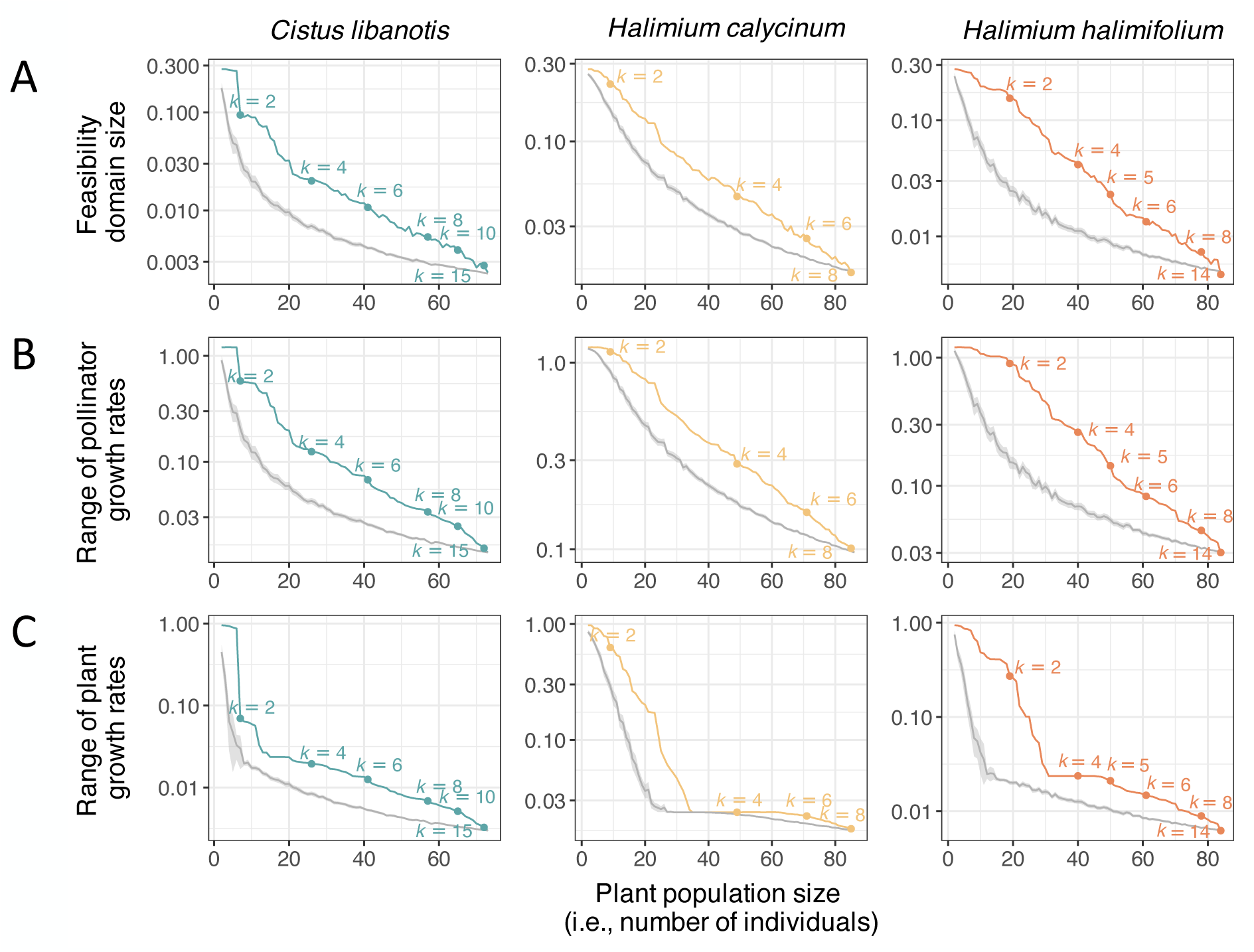
Effects of a higher proportion of specialized plant individuals in pollinator use, compared to mixtures of specialized and generalized plant individuals, on the persistence of plant populations’ mutualistic assemblages. For the three focal plant species (different colors), gray lines represent the size of the feasibility domain of the entire assemblage (A), and the range of feasible growth rates of the plant species (B) and of the pollinator community (C) over an increasing population size by including plant individuals regardless of their degree. Colored lines represent the size of the feasibility domain of the entire assemblage (A), and the range of feasible growth rates of the plant species (B) and of the pollinator community (C) over an increasing plant population size by incorporating plant individuals from lower to a higher degree (i.e., from more specialized to more generalized individuals). Plant individuals with the same degree were included in order of decreasing specificity (i.e., decreasing coefficient of variation of interactions). Colored dots represent the degree (*k*) of plant individuals being included to increase population size. Differences in the size of feasibility domain and in the ranges of plant and pollinator feasible growth rates between the colored and gray lines for a given population size correspond to the effects of having a higher proportion of specialized individuals in the focal species, compared to a mixture of specialized and generalized individuals. Note that the lines corresponding to a higher proportion of specialized individuals (colored) are located above the confidence interval (95%) of the random mixtures of specialized and generalized (gray) across all population sizes.

### Effects of within-population individual specialization across the plant community

Beyond its effects at the population level, the presence of more specialized individuals within specific plant species had implications at the community level by reshaping the vulnerability of these focal plant species as well as other species in the community. In contrast to the population-level results, having a higher proportion of specialized plant individuals did not necessarily create positive effects on the persistence of the focal population when it is embedded in a broader plant community, and therefore, subject to heterospecific plant competition. In fact, we found that in two of the focal plant species, having a higher proportion of specialized individuals increased the vulnerability of these focal species while it decreased the vulnerability of any other plant species in the community (Fig. 5). For the other focal plant species, we did not find strong effects of the presence of having more specialized individuals on its own vulnerability and only found weak contrasting effects on the vulnerability of co-occurring species. This is likely due to specific features of this focal species, such as the low levels of mutualistic benefits received (see Figure S8) or the limited flowering overlap with other plant species, which may decrease the magnitude of these cascading effects across the community.

**Figure 5:**
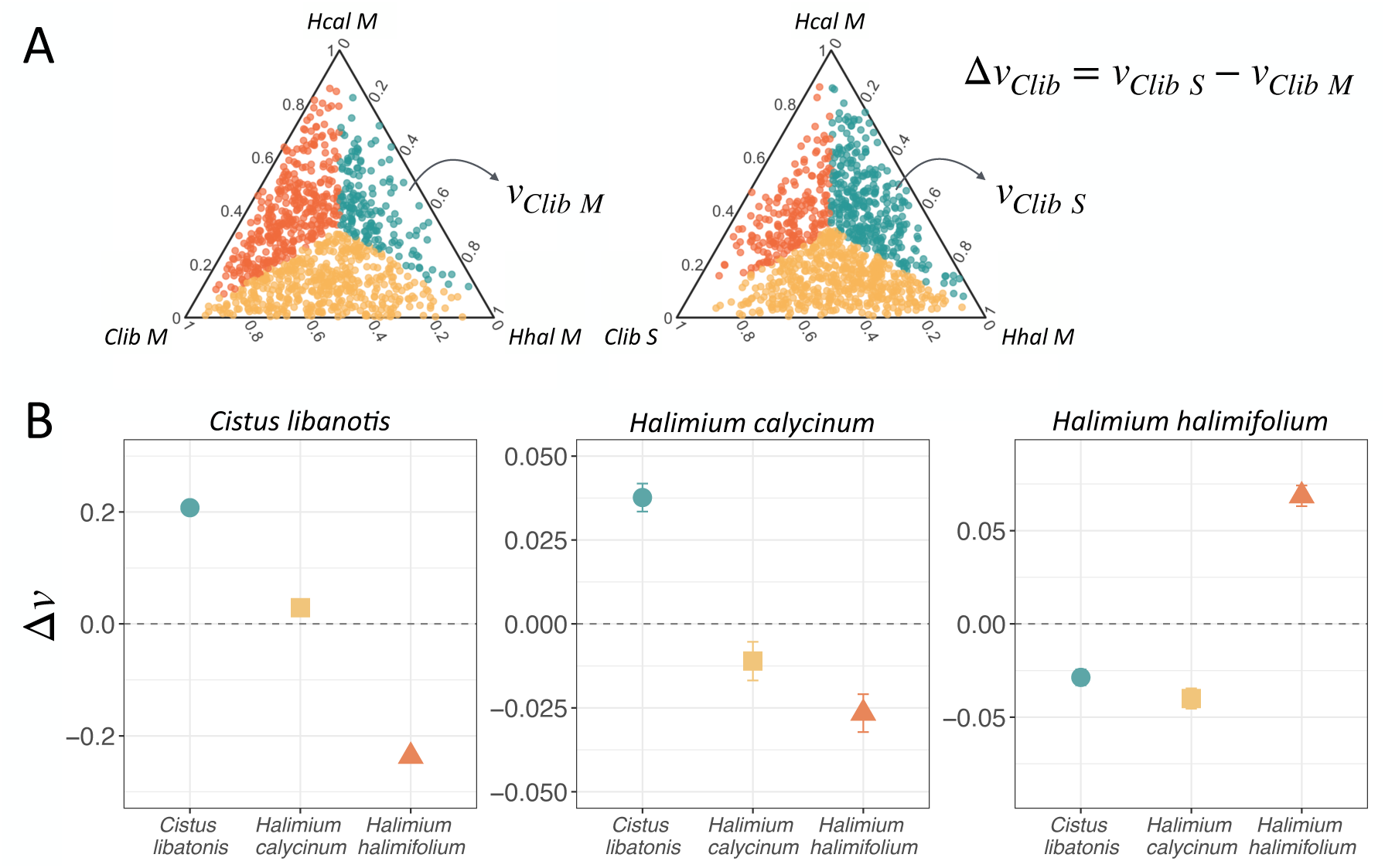
(A) Conceptual representation of the procedure used to measure the effects of having a higher proportion of specialized individuals within a focal plant species (different colors) on the vulnerability (v) of itself and co-occurring species, compared to having mixtures of generalized and specialized individuals in this focal species. For this conceptual representation, *Cistus libanotis* is used as the focal species. Dots represent vectors of feasible densities of the three plant species, which are colored based on the plant species with the lowest normalized density (i.e., highest vulnerability), represented in the axes of the ternary diagrams. The scenario on the left represents a focal population composed of a mixture of specialized and generalized individuals (*Clib M*), while the scenario on the right represents a focal population composed of a higher proportion of specialized individuals (*Clib S*). For each plant species, the greater the number of colored dots in the *S* scenario compared to the *M* scenario, the higher the vulnerability of that plant species when the focal plant population is composed of a higher proportion of specialized individuals. Note that here we focus only on the vulnerability of the plant side of the plant–pollinator community. (B) Effects of a higher proportion of specialized individuals within three focal plant species (indicated at the top of each panel) on the vulnerability (v) of itself and other species. A positive or negative Δv indicates that a higher proportion of specialized individuals of a focal plant species increases or decreases, respectively, the vulnerability of itself and co-occurring plant species, denoted on the *x*-axis and represented with different colors and symbols. Error bars around dots represent variation (95% CI) between plant populations composed of random mixtures of N=40 individuals (i.e., including individuals with high and low degree within the population). In each focal species, a higher proportion of specialized plant individuals was created using populations composed of the N=40 most specialized (lowest degree) individuals.

## DISCUSSION

Variation within populations has been identified as important as variation across species in ecological communities (Albert *et al*., 2010; Siefert *et al*., 2015), and thus, may potentially impact the functioning and stability of ecological systems (Bolnick *et al*., 2011; Violle *et al*., 2012). While widely recognized, the effects of intraspecific variation in shaping the dynamics of communities, especially those involving mutualistic partners, are virtually unknown. Downscaling to the level of individuals to understand ecological processes at this scale poses logistical and tractability challenges for empiricists and theoreticians. However, new sampling techniques now enable the collection of individual-level data with unprecedented accuracy (e.g., Besson *et al*., 2022; Corcoran *et al*., 2021; Droissart *et al*., 2021; Lines *et al*., 2022; Nathan *et al*., 2022; Steen, 2017), and recent developments have expanded the analytical toolbox to deal with these data (e.g., Coblentz *et al*., 2017; McCaslin *et al*., 2021; Sibly *et al*., 2013). The theoretical framework we propose here offers a pathway for further exploration of how intraspecific variation in interaction patterns shapes fitness variation, population dynamics, and the lasting effects of this source of variation within communities.

By leveraging the proposed mathematical framework, we demonstrated how intraspecific variation can shape species persistence in local communities, integrating theoretical predictions with insights from a detailed empirical case study. This comprehensive approach underscores the importance of individual-level variation in community dynamics and reveals key mechanisms promoting diversity. In our study system, we found that, when considering a single plant species without heterospecific plant competitors, plant populations composed of a higher proportion of specialized individuals promoted the persistence of the plant species they belong to and the pollinator community, compared to a plant population with heterogeneous mixtures of individuals. However, we found that these positive effects of having many specialized individuals largely vanished when plant species are embedded in a broader community of co-occurring plant species. In such contexts, having more heterogeneous mixtures of plant individuals may be more beneficial. As expected, focal plant species differed in the magnitude of the effects they exerted across their community, which was likely due to differences in the mutualistic benefits they received. Overall, our results suggest that the presence of heterogeneous plant populations, whose individuals vary in the level of specialization in resource use, may promote diversity in mutualistic communities because such individual variation changes the overall patterns of direct and indirect pairwise interactions between species. Our findings are particularly critical in the context of global change scenarios, as populations are expected to experience significant declines in size, become more homogeneous in traits, and exhibit a reduced level of generalization in resource use (Sala *et al*., 2000; Tylianakis *et al*., 2008).

Intraspecific variation in traits and ecological interactions is a significant driver of fitness variation within mutualistic communities (Arroyo-Correa *et al*., 2021; Gómez & Perfectti, 2012). Our approach incorporates variation in traits, mutualistic partners, and visitation rates across plant individuals, all of which determine the fitness benefits received. An alternative approach that relies on a species-level value for each of these features, averaged across individuals, to estimate mutualistic benefits could result in inaccurate feasibility estimations at both population and community levels. In other words, a species-level value to parameterize mutualistic benefits calculated from averaged interaction strengths and traits exhibited by individuals is unlikely to represent the mutualistic benefits received by any particular individual within a given population. Consequently, using interaction and trait data averaged across individuals to calculate the fitness benefits received by a given species from its mutualistic partners may lead to biased predictions of species persistence. Moreover, our findings challenge conventional assumptions about the distribution of mutualistic benefits based solely on the number of mutualistic partners per species (e.g., Rohr *et al*., 2014; Saavedra *et al*., 2013, 2016b). As we have shown, the benefits received by mutualistic partners cannot be accurately summarised by only considering the number of mutualistic partners at the species level, as they are ultimately defined by multiple factors that come into play at the level of individuals. We anticipate that species-level, aggregated analyses (e.g., based on average estimates of generalization/specialization in resource use) may underestimate the potential of species to persist in real communities. Therefore, our results suggest a more nuanced interaction outcome that warrants further exploration in order to effectively represent fitness variation in natural populations and capture this kind of variation in mutualistic communities’ dynamic models.

While saturation effects in the benefits individuals receive can be important in some mutualistic systems, we opted for a linear functional response to incorporate mutualistic benefits in the proposed mathematical framework in order to isolate the effects of intraspecific variation in mutualistic interaction patterns on community dynamics. Including saturation would have introduced additional complexity in estimating the size of the feasibility domain (Cenci & Saavedra, 2018), potentially obscuring the specific contributions of this variation. Importantly, we also did not observe evidence of saturation in the mutualistic benefits received by individuals of any plant species considered in our empirical case study (i.e., we did not detect pollination limitation). In fact, pollen limitation is reported to be common in natural systems (Knight *et al*., 2005), suggesting most systems are not near saturation. Thus, we considered a linear response to be a reasonable assumption for this specific context. Nonetheless, we acknowledge that mutualistic benefits may decrease at high interaction rates in some systems, making it important to account for saturation effects in certain contexts. Exploring these non-linearities could provide a valuable extension to the framework, particularly in systems where saturation plays a more significant role.

By considering various forms of variation within populations, we are able to identify thresholds of minimum amount of variation that would need to be preserved to guarantee the conservation of stable and biodiverse local communities (Anderegg *et al*., 2013; Brose *et al*., 2012; Clark, 2010). For instance, if disturbance effects simplify populations by size-dependent mortality events (Bennett *et al*., 2015; Gora & Esquivel-Muelbert, 2021), one can assess how this kind of perturbation may impact among-individual variance and its effects on the feasibility of the entire communities in which these populations are embedded. Moreover, our framework enables the identification of keystone individuals, as well as their attributes, that exert disproportionately large effects on population and community dynamics (Garrido-de León *et al*., 2024; LaBarge *et al*., 2024; Modlmeier *et al*., 2014). Beyond considering the role of keystone species (Cagua *et al*., 2019; Jordán, 2009; Mello *et al*., 2015; Timóteo *et al*., 2023), understanding the contributions of key individuals allows us to better predict the resilience of mutualistic communities to environmental perturbations and human-induced disturbances. Only by downscaling from species to the most basic interacting elements (i.e., individuals) when assessing the dynamics of natural systems can we uncover the fine-scale mechanisms that foster species persistence, which would otherwise remain obscured (Cantwell-Jones *et al*., 2024). Therefore, an individual-based approach may improve our predictive power regarding the structure and functioning of ecological systems under the effects of global change. This improved understanding would enable the development of conservation strategies that support the persistence of functional co-occurring populations within natural communities, such as planting trees of varying age classes or preserving size diversity and/or size hierarchies within populations (Des Roches *et al*., 2017; Hughes *et al*., 1997).

## FUTURE DIRECTIONS

Given the critical role of intraspecific variation in shaping mutualistic communities, future research should prioritize unveiling the full impact of this variation within species across different ecological contexts and spatial scales. One promising direction involves extending the current framework to study the effects of intraspecific variation in a broader range of species and ecosystems, particularly under different environmental conditions. Upcoming research applying this framework to assess how intraspecific variation promotes the persistence of communities in disturbed ecosystems, such as those especially affected by climate change, would provide valuable insights into the mechanisms that support biodiversity.

In our empirical case study, we focused on assessing intraspecific variation in mutualistic benefits from a phytocentric perspective. In doing so, we ignored the effects of co-variation that could arise when also considering the fact that animal populations can be potentially composed of individuals differing in traits and ecological interactions (Bascompte & Jordano, 2007). One main limitation of considering both partners variability lies in the logistical challenges associated with gathering empirical data on the mutualistic benefits received by animal partners (Vázquez *et al*., 2012), particularly when they are invertebrates such as insect pollinators. Future research efforts that transcend these sampling difficulties could gain from incorporating perspectives that focus on both plants and animals to comprehensively explore the overall picture of the influence of intraspecific variation on mutualistic communities. If these fine-scale data can be collected in the field, our dynamic model accounting for intraspecific variation is ready to be parameterized to gain even deeper insights into the ecological processes that shape mutualistic communities.

Our main focus here was to specifically examine the role of variation in mutualistic interaction patterns among individuals within a single overarching population. To achieve this narrowing of focus, we assumed that all individual types share the same growth rate, allowing us to treat them as part of a single, “cohesive” population. Nevertheless, we acknowledge that this parameter is likely to show some degree of variation within real-world populations. Therefore, incorporating such variation presents an exciting avenue for future research, offering the potential to complement our findings by shedding light on the relative contributions of these two sources of intraspecific variation to community dynamics. Moreover, our theoretical framework also has broader applications beyond testing the effects of intraspecific variation in mutualistic interactions, as it could be also be extended to other types of species interactions or organizational levels within ecological systems for which Lotka–Volterra models are useful to describe dynamics (Boucher, 1985; Pascual-García & Bastolla, 2017). Mutualism rarely occurs in isolation, but in combination with other types of interactions (e.g., antagonistic) so expanding this framework to multiple interaction types can be useful to test the extent to which within-species variation in various types of interactions can shape communities. We also offer a versatile tool for exploring dynamics at various scales, from genotypes within populations to communities within metacommunities and beyond. Thus, the flexibility of this tool allows us to explore various timely questions in ecology involving any kind of subsetting structure across multiple scales (Chase *et al*., 2018). For instance, our framework facilitates delving into the temporal stability of genetically-structured populations, shedding light on the resilience and persistence of such populations over time (Booy *et al*., 2000). The development of more sophisticated models that integrate intraspecific variation with other sources of ecological complexity, such as multi-trophic interactions, temporal dynamics, and spatial heterogeneity, represents a promising opportunity for further exploration.

Advancing methodological techniques to collect and capture fine-scale individual-level data across a broader range of species, particularly those that are difficult to study due to their intrinsic characteristics, will be essential. Leveraging new technologies, such as remote sensing, DNA barcoding, and automated tracking systems, can enhance our ability to collect high-resolution data on individual interactions and traits in natural settings (Besson *et al*., 2022; Corcoran *et al*., 2021; Kellner *et al*., 2019; Koger *et al*., 2023; Kress *et al*., 2015; Lines *et al*., 2022). These data could then be used to refine and parameterize models, improving the accuracy of predictions related to community dynamics and species persistence. By combining detailed empirical data with theory and mathematical models, we open the door to deeply understanding ecological processes underpinning diversity patterns.

In summary, future research directions should aim to broaden the scope of intraspecific variation studies across ecological contexts, refine theoretical models, enhance empirical data collection, and adopt interdisciplinary approaches to better understand the complex interplay between individual variation and the maintenance of diversity. These efforts will enhance our predictive capabilities, which are crucial for addressing challenges posed by global change and mitigating its impacts on biodiversity and ecosystem services.

## Acknowledgements

We thank the logistic support and facilities provided by ICTS-Doñana and Doñana National Park for onsite access authorizations during the fieldwork. B.A.-C. received funding from the Ministry of Universities of the Spanish Government (FPU19/02552 and EST23/00036) and a project grant from the Spanish Association of Terrestrial Ecology (AEET). B.A-C. and P.J. were supported by grants CGL2017-82847-P and PID2022-136812NB-I00, funded by MCIN/AEI/10.13039/ 501100011033 and by the “European Union NextGenerationEU/PRTR”, grant P20-00736 from Junta de Andalucía, and a LifeWatch ERIC-SUMHAL project (LIFEWATCH-2019-09-CSIC-13), with FEDER-EU funding. E.F.C. acknowledges the support from the University of Canterbury Doctoral Scholarship, the University of Canterbury Meadow Mushrooms Postgraduate Scholarship, and a New Zealand International Doctoral Research Scholarship. D.B.S. acknowledges the support of a Rutherford Discovery Fellowship (RDF-13-UOC-003), administered by the Royal Society of New Zealand Te Apārangi.

## Box 1.

Empirical estimation of mutualistic benefits.

To parameterize the mutualistic benefits provided by each pollinator species, the first step is to specifically estimate the contribution of each pollinator species to a plant individual’s fitness of a given plant species as the number of seeds produced per flower as a result of a single visit by each pollinator species. First, as the three focal plant species (*Cistus libanotis*, *Halimium calycinum*, *Halimium halimifolium*) produce fruits without a fixed maximum number of seeds, the number of seeds produced per fruit recorded in each plant individual can be modeled as a Poisson outcome with:

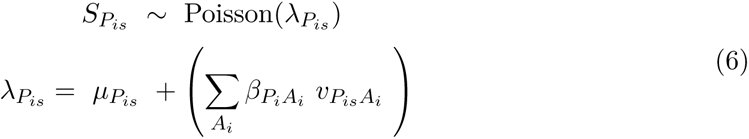

where *SP_is_*is the observed number of seeds produced in an individual fruit of plant individual *s* belonging to plant species *i*, *⋋P_is_* is the mean expected intensity, *µP_is_* represents the intercept for plant individual *s* belonging to plant species *i*, *βP_is_A_i_* the coefficient of pollinator species *i* on plant species *i*, and *vP_is_A_i_* the frequency of interactions per flower between pollinator species *i* and plant individual *s* of plant species *i* along the entire flowering season (estimated using the visitation data we recorded in the field; Appendix B).

Second, since a flower can only produce seeds if it has previously set fruit, we also need to estimate the contribution of each pollinator species to the probability of a flower setting fruit. The fruit production per individual flower in each plant individual was recorded as either the presence or absence of fruit, and therefore can be treated as a Bernoulli outcome:

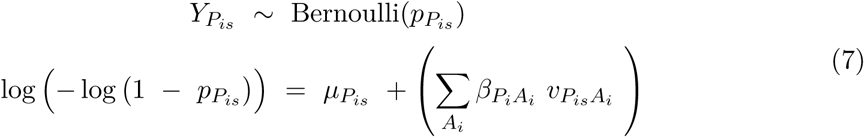

where *YP_is_*is the response variable denoting the presence or absence of a fruit on an individual flower of plant individual *s* belonging to plant species *i*, *pP_is_*is the probability of the presence of a fruit, and all other parameters are defined above.

We integrated both fitness proxies—the probability of a flower setting fruit and the expected number of seeds per fruit—into a single model using a joint likelihood (Appendix C). The joint likelihood method simultaneously fits a likelihood to both data sources (fruit and seed production) while accounting for different data types but using equivalent parameter values for both (Ahmad Suhaimi *et al*., 2021; Simmonds *et al*., 2020). This is the reason why we used a cloglog link in the Bernoulli model (Eq. 7) to connect *pP_is_* to the log intensity (*⋋P_is_*) of the Poisson model (Eq. 6), so that *pP_is_ ≡* 1 *− e^−⋋P^_is_* (Royle *et al*., 2009). As a result, the parameters *βP_is_A_i_* are directly shared between both models, and we can simultaneously estimate the contribution of pollinator species *i* to both the probability of a flower setting fruit and the number of seeds produced per fruit in plant species *i* (Appendix C). Note that this equation implies that the probability of a flower not producing a fruit is equal to the probability of a fruit from that flower not producing any seeds.

With this statistical procedure, we estimated the contribution of each pollinator species to seed production per flower in a given plant species (*βP_i_A_i_*). In total, we inferred a separate *β* parameter for each pollinator species visiting each plant species, totaling 55 in *Cistus libanotis*, 33 in *Halimium calycinum* and 38 in *Halimium halimifolium* (Appendix C). These pollinator species’ contributions were then used to determine the specific mutualistic benefits received by plant individuals from each pollinator species, expressed as *ϒP_is_A_i_* in matrix *M* of the mathematical framework for our specific case (Eq. 5). We estimated the mutualistic benefit of pollinator species *i* (*A_i_*) to plant individual type *s* belonging to plant species *i* (*P_is_*) as:

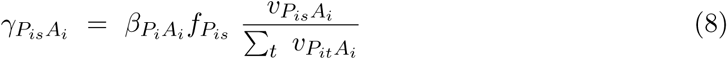

where *fP_is_* is the total number of flowers produced by plant individual type *s* belonging to plant species *i*, and *vP_is_A_i_* is the frequency of visits per flower received by plant individual type *s* of plant species *i* from pollinator species *i* across the entire flowering season, which is normalized across all individuals of plant species *i*. Therefore, intraspecific variation in mutualistic benefits received by plant individuals from pollinator species (i.e., *ϒP_is_A_i_*) within a plant population is modulated by three components: pollinator assemblage and their associated contribution to fitness (*βP_i_A_i_*), visitation rates (*vP_is_A_i_*) and flower production across the flowering season (*fPis*). Note that *ϒP_is_A_i_* = 0 in the case that pollinator species *Ai* did not establish interactions with plant individual type *P*_is_ (*vP_is_A_i_* = 0).

## Box 2.

Theoretical insights into the role of intraspecific variation in the feasibility of simplified mutualistic communities.

To explore the theoretical implications of the presence of intraspecific variation, we studied a simplified system consisting of a plant population comprising two individual types, *P*_11_ and *P*_12_, both of which might potentially receive visits from two pollinator species, *A*_1_ and *A*_2_ (precisely as given by Eq. 5). We examined different cases in which plant individual types may or may not differ in interaction patterns, including a case where both types are specialized (‘fully specialized’), both types are generalized (‘fully generalized’), and cases where we have mixtures of specialized and generalized individuals (Fig. 1). Consistent with our empirical data, we allowed the pollinator species to differ in abundance (*A*_1_ *< A*_2_), and the less abundant pollinator species contributes more to plant fitness per visit (*βP*_1_*A*_1_ *> βP*_1_*A*_2_) (Fig. 1A). We set *A*_1_=0.4, *A*_2_=0.6, *βP*_1_*A*_1_ =0.4 and *βP*_1_*A*_2_ =0.2, with interaction coefficients parameterized as described in Appendix E.

For each of the cases included in this theoretical testing, we calculated the size of the feasibility domain of a system in which the plant population is composed of each of the single individual types by itself (without intraspecific variation) and contrasted this to when the population is composed of both types (with intraspecific variation). Treating the data in this way thus allows us to remove and isolate the effect of intraspecific variation (Fig. 1B). We started with four specific cases because they represent the most extreme scenarios. However, in real-world systems, populations with relatively generalized individuals can effectively function as a mix of generalized and specialized individuals due to substantial differences in visitation rates. To account for this possibility, we also calculated the size of the feasibility domain across a range of variation in visitation rates between the two plant individual types (Fig. 1C), spanning these extreme cases as well as other combinations with different visitation rates.

Just as feasibility of multi-species communities results from a balance of direct and indirect effects *between* species (Bastolla *et al*., 2009; Rohr *et al*., 2014; Saavedra *et al*., 2016a), feasibility of a population comprising mixtures of individual types also depends on a balance of direct and indirect effects *within* species. As a result, the feasibility domain for a population with intraspecific variation in interactions will not simply be the intersection between the feasibility domains of each of that population’s constituent individual types (Fig. 1B).

Importantly, we found that a system with two distinct individual types can only be feasible within a very narrow range of conditions (Fig. 1C). Indeed, just as two co-occurring species require sufficiently different interactions to avoid competitive exclusion (MacArthur & Levins, 1967), there is an upper bound on the amount of variation that two co-occurring individual types belonging to the same species can show before one type ultimately “out-competes” the other. This upper bound is determined by a balance between differences in pollinators’ abundances and contributions, which dictates the maximum level of specialization of plant individual types that cannot be surpassed for the system to be feasible (Fig. S6).

