## Appendix for "Integrating intraspecific variation into population dynamics reveals how interacting species persist in mutualistic communities"

### A Imposing constraints on feasibility analyses

As described in the main text, at the core of our analyses of the feasibility domain is the equation that must hold at any equilibria (feasible or otherwise) given by

$$\overbrace{\begin{bmatrix} r_{P_{is}} \\ r_{P_{it}} \\ r_{A_i} \\ r_{A_j} \end{bmatrix}}^{\mathbf{r}} = \overbrace{\begin{bmatrix} \alpha_{P_{is}P_{is}} & \alpha_{P_{is}P_{it}} & -\gamma_{P_{is}A_i} & -\gamma_{P_{is}A_j} \\ \alpha_{P_{it}P_{is}} & \alpha_{P_{it}P_{it}} & -\gamma_{P_{it}A_i} & -\gamma_{P_{it}A_j} \\ -\gamma_{A_iP_{is}} & -\gamma_{A_iP_j} & \alpha_{A_iA_i} & \alpha_{A_iA_j} \\ -\gamma_{A_jP_{is}} & -\gamma_{A_jP_j} & \alpha_{A_jA_i} & \alpha_{A_jA_j} \end{bmatrix}}^{M} \overbrace{\begin{bmatrix} P^*_{is} \\ P^*_{it} \\ A^*_i \\ A^*_j \end{bmatrix}}^{\mathbf{N}^*}. \tag{S1}$$

Most conventionally, the only condition imposed on the vectors $\mathbf{r}$ is that they provide a feasible equilibrium for all densities in the vector $\mathbf{N}^*$. One can directly estimate the size of the feasibility domain dictated by the matrix $M$ as the fraction of randomly sampled growth-rate vectors $\mathbf{r}$ on the unit sphere (i.e., for which $(\sum_{x,y}(r_{P_{xy}})^2) + (\sum_x(r_{A_x})^2) = 1$) whose corresponding equilibrium densities $\mathbf{N}^*$ are all positive. For the community whose equilibrium is determined by Eq. S1, each of these candidate growth-rate vectors falls on the surface of a 4-dimensional hypersphere. We describe here how our two additional types of constraints change the underlying properties of the vector $\mathbf{r}$—in particular its number of free parameters—necessary for this Monte Carlo estimation process.

##### Pollinator density constraints

In one case, we are interested in feasible equilibria for which pollinator species' relative abundances are known. If, for example, we require $A^*_i = \pi_{A_i} A^*$ where $A^* = A^*_i + A^*_j$, then we could rewrite

Eq. S1 as

$$\begin{bmatrix} r_{P_{is}} \\ r_{P_{it}} \\ r_{A_i} \\ r_{A_j} \end{bmatrix} = \begin{bmatrix} \alpha_{P_{is}P_{is}} & \alpha_{P_{is}P_{it}} & -\gamma_{P_{is}A_i} & -\gamma_{P_{is}A_j} \\ \alpha_{P_{it}P_{is}} & \alpha_{P_{it}P_{it}} & -\gamma_{P_{it}A_i} & -\gamma_{P_{it}A_j} \\ -\gamma_{A_iP_{is}} & -\gamma_{A_iP_j} & \alpha_{A_iA_i} & \alpha_{A_iA_j} \\ -\gamma_{A_jP_{is}} & -\gamma_{A_jP_j} & \alpha_{A_jA_i} & \alpha_{A_jA_j} \end{bmatrix} \overbrace{\begin{bmatrix} 1 & 0 & 0 \\ 0 & 1 & 0 \\ 0 & 0 & \pi_{A_i} \\ 0 & 0 & 1-\pi_{A_i} \end{bmatrix}}^{\Pi} \overbrace{\begin{bmatrix} P^*_{it} \\ P^*_{is} \\ A^* \end{bmatrix}}^{\tilde{\mathbf{N}}^*}, \tag{S2}$$

where the new matrix $\Pi$ that appears after $M$ captures the explicit density constraints between species. Note that the values in the corresponding column of the matrix $\Pi$ must sum to one. As a consequence, for any potential equilibrium given by the three densities in the reduced vector $\tilde{\mathbf{N}}^* = \{P^*_{is}, P^*_{it}, A^*\}$, we could immediately determine all four growth rates consistent with that equilibrium. We can see how this process results mathematically in a reduction of the number of free growth rates by instead rewriting Eq. S1 as

$$\overbrace{\begin{bmatrix} r_{P_{is}} \\ r_{P_{it}} \\ r_{A_i} \\ 0 \end{bmatrix}}^{\tilde{\mathbf{r}}} = \overbrace{\begin{bmatrix} \alpha_{P_{is}P_{is}} & \alpha_{P_{is}P_{it}} & -\gamma_{P_{is}A_i} & -\gamma_{P_{is}A_j} \\ \alpha_{P_{it}P_{is}} & \alpha_{P_{it}P_{it}} & -\gamma_{P_{it}A_i} & -\gamma_{P_{it}A_j} \\ -\gamma_{A_iP_{is}} & -\gamma_{A_iP_j} & \alpha_{A_iA_i} & \alpha_{A_iA_j} \\ 0 & 0 & 1-\pi_{A_i} & -\pi_{A_i} \end{bmatrix}}^{\tilde{M}} \begin{bmatrix} P^*_{is} \\ P^*_{it} \\ A^*_i \\ A^*_j \end{bmatrix}, \tag{S3}$$

where the change to the last element in the growth-rate vector and last row of the matrix $\tilde{M}$ corresponds precisely to the density constraint between $A_i$ and $A_j$. The fact that $r_{A_j}$ no longer appears anywhere in this rewritten equation explicitly dictates that it is no longer a free parameter. Indeed, for a candidate growth-rate vector $\tilde{\mathbf{r}}$ with a zero as its fourth element, we can determine its full set of four equilibrium densities with $\tilde{M}^{-1}\,\tilde{\mathbf{r}}$. As a result, candidate growth-rate vectors required for the Monte Carlo estimation in this scenario fall on a 3-dimensional hypersphere.

In our specific empirical case study, pollinator abundance was defined as $\pi_{A_i} = \frac{\sum_{P_{is}} m_{P_{is}A_i}}{\sum_{P_{is},A_i} m_{P_{is}A_i}}$ in Eq. S2, where $m_{P_{is}A_i}$ is the number of visits per minute by pollinator species $i$ to plant individual $s$ of plant species $i$.

#### Plant type growth-rate constraints

In the second case, we are interested in feasible equilibria for which some species' growth rates are exactly equal. If, for example, we require that $r_{P_{it}} = r_{P_{is}}$, then we could rewrite Eq. S1 as

$$\overbrace{\begin{bmatrix} r_{P_{is}} \\ 0 \\ r_{A_i} \\ r_{A_j} \end{bmatrix}}^{\tilde{r}} = \overbrace{\begin{bmatrix} \alpha_{P_{is}P_{is}} & \alpha_{P_{is}P_{it}} & -\gamma_{P_{is}A_i} & -\gamma_{P_{is}A_j} \\ \alpha_{P_{it}P_{is}} - \alpha_{P_{is}P_{is}} & \alpha_{P_{it}P_{it}} - \alpha_{P_{is}P_{it}} & -\gamma_{P_{it}A_i} + \gamma_{P_{is}A_i} & -\gamma_{P_{it}A_j} + \gamma_{P_{is}A_j} \\ -\gamma_{A_iP_{is}} & -\gamma_{A_iP_j} & \alpha_{A_iA_i} & \alpha_{A_iA_j} \\ -\gamma_{A_jP_{is}} & -\gamma_{A_jP_j} & \alpha_{A_jA_i} & \alpha_{A_jA_j} \end{bmatrix}}^{\tilde{M}} \begin{bmatrix} P^*_{is} \\ P^*_{it} \\ A^*_i \\ A^*_j \end{bmatrix}, \quad \text{(S4)}$$

where the changes in the growth-rate vector to create $\tilde{r}$ and in the interaction matrix to create $\tilde{M}$ correspond to the aforementioned growth-rate constraints. As above, the fact that $r_{P_{it}}$ no longer appears in this rewritten equation implies that it is no longer a free parameter. That is, if we only know the values for $r_{P_{is}}$, $r_{A_i}$ and $r_{A_j}$, we could immediately solve this equation for the four densities in $\mathbf{N}^*$. As a result, candidate growth-rate vectors required for the Monte Carlo estimation in this scenario fall on a 3-dimensional hypersphere.

To perform this operation "in reverse", we need to recognize that the growth-rate constraint captured by the second row in Eq. S4 implies that $P^*_{it}$ can be determined from $P^*_{is}$, $A^*_i$, and $A^*_j$ as given by

$$\overbrace{\begin{bmatrix}\alpha_{P_{is}P_{it}} - \alpha_{P_{it}P_{it}}\end{bmatrix}}^{L} \begin{bmatrix}P_{it}^*\end{bmatrix} = \overbrace{\begin{bmatrix}\alpha_{P_{it}P_{is}} - \alpha_{P_{is}P_{is}} & -\gamma_{P_{it}A_i} + \gamma_{P_{is}A_i} & -\gamma_{P_{it}A_j} + \gamma_{P_{is}A_j}\end{bmatrix}}^{R} \overbrace{\begin{bmatrix}P_{is}^* \\ A_i^* \\ A_j^*\end{bmatrix}}^{\tilde{\mathbf{N}}^*} . \quad \text{(S5)}$$

Therefore, we could again rewrite Eq. S1 as

$$\begin{bmatrix} r_{P_{is}} \\ r_{P_{it}} \\ r_{A_i} \\ r_{A_j} \end{bmatrix} = \begin{bmatrix} \alpha_{P_{is}P_{is}} & \alpha_{P_{is}P_{it}} & -\gamma_{P_{is}A_i} & -\gamma_{P_{is}A_j} \\ \alpha_{P_{it}P_{is}} & \alpha_{P_{it}P_{it}} & -\gamma_{P_{it}A_i} & -\gamma_{P_{it}A_j} \\ -\gamma_{A_iP_{is}} & -\gamma_{A_iP_j} & \alpha_{A_iA_i} & \alpha_{A_iA_j} \\ -\gamma_{A_jP_{is}} & -\gamma_{A_jP_j} & \alpha_{A_jA_i} & \alpha_{A_jA_j} \end{bmatrix} \overbrace{\begin{bmatrix} 1 & 0 & 0 \\ & L^{-1}R & \\ 0 & 1 & 0 \\ 0 & 0 & 1 \end{bmatrix}}^{\rho} \overbrace{\begin{bmatrix} P_{is}^* \\ A_i^* \\ A_j^* \end{bmatrix}}^{\tilde{\mathbf{N}}^*}, \quad \text{(S6)}$$

where the $1 \times 3$ matrix $L^{-1}R$ provides the second row of the matrix $\rho$. That is, for any potential equilibrium given by the three densities in the vector $\tilde{\mathbf{N}}^* = \left\{P_{is}^*, A_i^*, A_j^*\right\}$, the resulting vector of four growth rates that are consistent with it will necessarily have $r_{P_{is}} = r_{P_{it}}$. Note, however, that just because a vector $\tilde{\mathbf{N}}^* = \left\{P_{is}^*, A_i^*, A_j^*\right\}$ is feasible (i.e., all values are $> 0$) does not imply that the vector $\mathbf{N}^* = \rho\,\tilde{\mathbf{N}}^*$ is also feasible because the resulting $P_{it}^*$ might be less than 0.

**Pollinator density and plant type growth-rate constraints**

In our analyses of matrices parameterized with empirical data, we simultaneously both of these types of additional constraints when estimating the size of the feasibility domain. A key benefit of doing so is that it reduces the dimension of the space of growth-rate vectors that must be randomly sampled when estimating the size of the feasibility domain (which also increases the probability of sampling feasible growth-rate vectors; Rohr *et al.*, 2014). Moreover and in contrast to a scenario where within-species plant types would be treated as separate “species”, each with their own unique growth rate, it also means that the dimension of the space of growth-rate vectors remains constant regardless of the number of individual types included within any given plant species. To use Monte Carlo approaches to estimate the size of the feasibility domain across our different scenarios (Song *et al.*, 2018; Song & Saavedra, 2018), in general our analyses require randomly sampled growth rate vectors on an $n$-dimensional hypersphere, where the dimension $n = n_{\text{plants}} + 1$ and $n_{\text{plants}}$ is the number of plant species. Note, however, that only one or none of our two types of constraints may be imposed on any given species or individual type at a time because of how each correspondingly reduces the number of free parameters.

### B Plant individuals' visitation rates

We estimated the contribution of pollinator species to plant individuals' fitness at the level of flowers ($\beta_{P_{is}A_i}$). Because data on plant fitness proxies were collected at the flower level on each plant individual, we used pollinator's visitation rates per flower, estimated using visitation data recorded in the field, to establish a link between plant individual's fitness and pollinator visitation when calculating these fitness contributions of pollinators. The visitation rate per flower of each pollinator species on each plant individual was estimated as:

$$v_{P_{is}A_i} = t_{P_{is}} \sum_q \frac{m_{P_{isq}A_i}}{f_{P_{isq}}} \tag{S7}$$

where $t_{P_{is}}$ is the total flowering time period (minutes) of plant individual $s$ of plant species $i$, $m_{P_{isq}A_i}$ is the number of visits per minute by pollinator species $i$ to plant individual $s$ of plant species $i$ in sampling event $q$, and $f_{P_{isq}}$ is the total number of active flowers in plant individual $s$ of plant species $i$ during sampling event $q$. By dividing the visitation rate per minute by the number of flowers active in the sampling event where the specific interaction was recorded, we aimed to calculate the rate of pollinator visitation *per flower* during that particular sampling event. This estimation allows for a more precise proxy of the efficiency or effectiveness of pollination resulting from visitation by potential pollinators to each plant individual.

#### C Inferring the contributions of pollinators to plant individuals' fitness

We estimated the values of $\beta_{P_iA_i}$ by fitting the model comprising Eqns. 6 and 7 with our observed fitness proxies ($S_{P_{is}}$ and $Y_{P_{is}}$) as the response variables and using observed visitation rates ($v_{P_{is}A_i}$) as predictor variables. We constrained all pollinator contributions to fitness to be positive ($\beta_{P_iA_i} \geq 0$). We also assumed that $\mu_{P_{is}} = 0$ for all plant individuals, implying that fruit and seed production is negligible in the absence of pollinator visits. This assumption is grounded in the biological characteristics of our three focal plant species, which depend on insect-mediated cross-pollination to complete fruit and seed production. Furthermore, our visitation sampling procedure comprehensively captured visitation rates to all plant individuals and reduced the likelihood of overlooking visitation rates (Arroyo-Correa *et al.*, 2023).

We performed these model fits separately for each focal plant species using the Automatic Differentiation Variational Inference approach (Kucukelbir *et al.*, 2017) provided by the `variational()` function in the `cmdstanr` package v0.7.1 (Gabry *et al.*, 2024) within `R` v4.3.1 (R Core Team, 2023). To improve convergence, we scaled observed visitation rates $v_{P_{is}A_i}$ by their standard deviation prior to fitting the model. We specified the following prior:

$$\tau_{P_iA_i} \sim \text{Normal}(0, 2) \tag{S8}$$

and converted these unconstrained parameters $\tau_{P_iA_i}$ to positively constrained parameters $\tilde{\beta}_{P_iA_i}$ with the transformation provided by the `log1p_exp()` function (i.e., $\tilde{\beta}_{P_iA_i} = \log\Big(1 + \exp(\tau_{P_iA_i})\Big)$. We used the "fullrank" algorithm, allowed $10^6$ iterations for convergence, and set initial values of $\tau_{P_iA_i} = 0$ for all unknown parameter values. We ensured that the `variational()` function reported successful model convergence for each focal plant species upon completion, and extracted a sample of 1000 posterior draws of the parameters $\tilde{\beta}_{P_iA_i}$.

We then "unscaled" the inferred parameter coefficients to create corresponding posterior distributions of the parameters $\beta_{P_iA_i}$ (i.e., $\beta_{P_iA_i} = \tilde{\beta}_{P_iA_i}/\sigma_{v_{P_{is}A_i}}$). We used the median of these posterior distributions of unscaled parameters $\beta_{P_iA_i}$ as the value of the fitness contribution of pollinator species $i$ to plant species $i$ (i.e., seeds produced per visit and per flower) required by our empirically parameterized population-dynamics model. For each focal species, we noticed that the inferred coefficients consisted of comparatively few pollinators making strong fitness contributions per visit whereas the majority of per visit fitness contributions were quite weak (Figs. S1–S3). Notably, this is consistent with observations in other systems and ecological contexts that empirically inferred interactions strengths tend to be sparse (Weiss-Lehman *et al.*, 2022; Lai *et al.*, 2024).

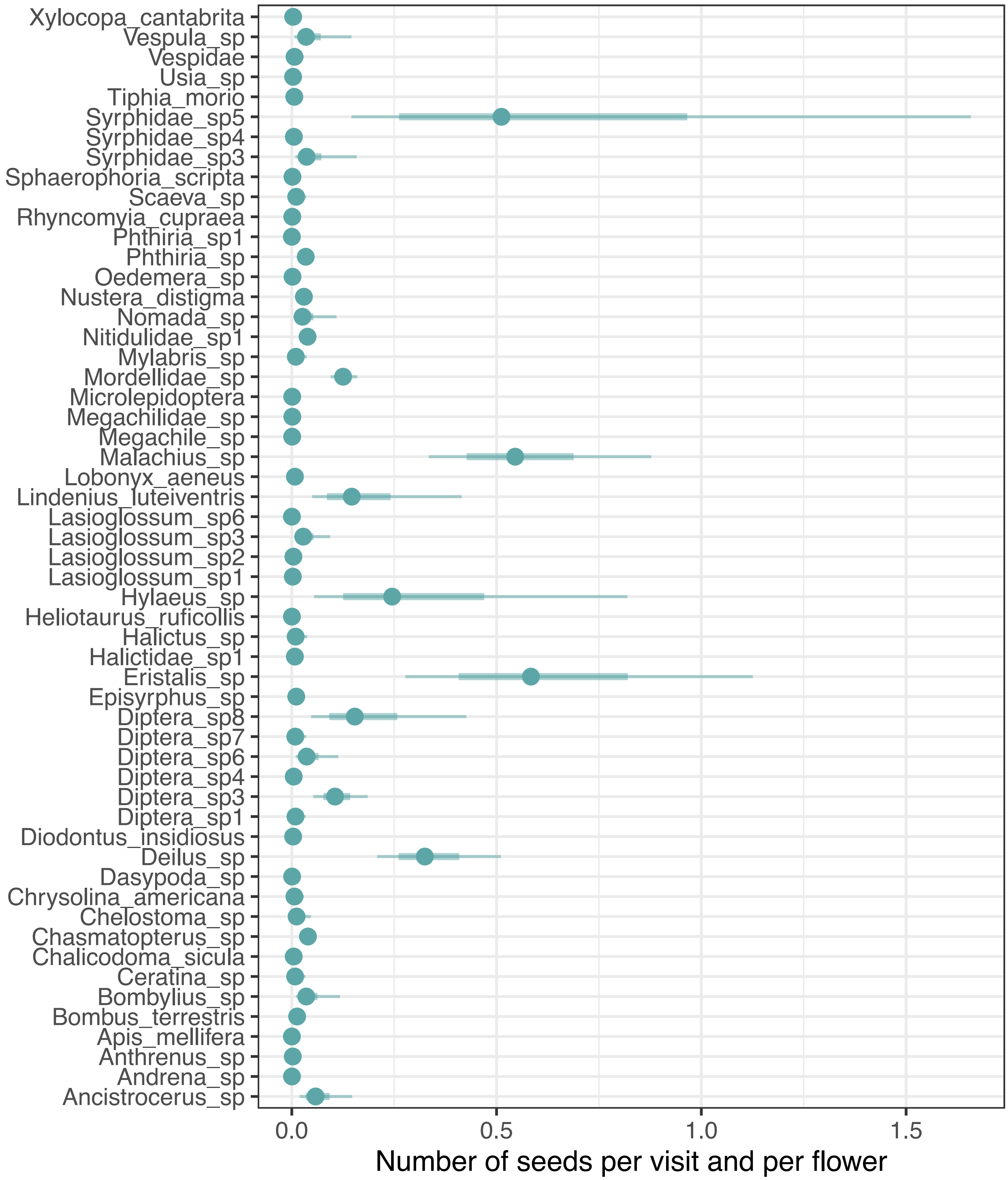


**Figure S1:** Credible intervals for parameters $\beta_{P_i A_i}$, which represent the value of the fitness contribution of pollinator species $i$ to plant species $i$ (i.e., seeds produced per visit and per flower), in this case *Cistus libanotis*. Points indicate the posterior medians, while thick segments and thinner outer lines represent 50% and 90% intervals, respectively.

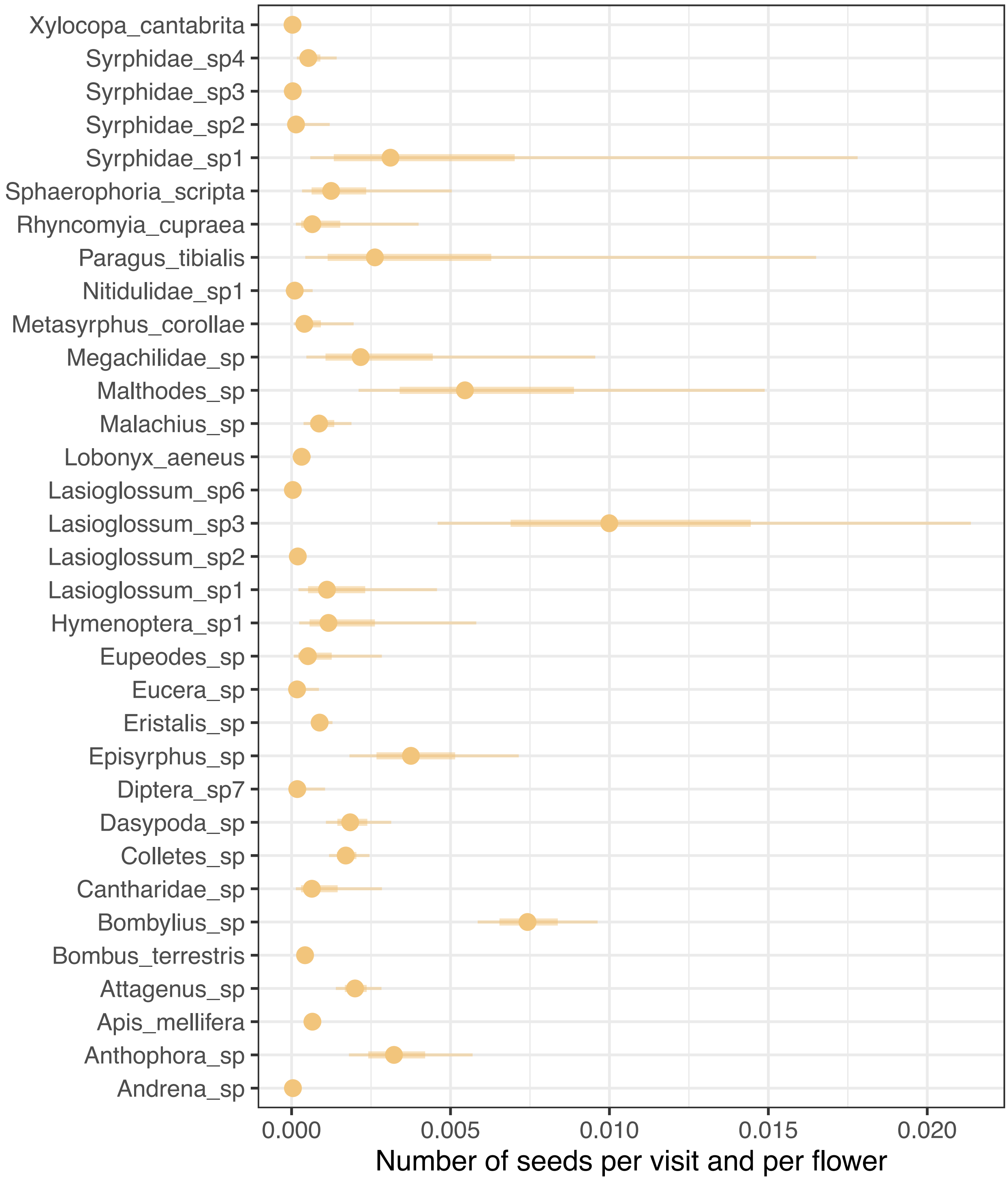


**Figure S2:** Credible intervals for parameters $\beta_{P_i A_i}$, which represent the value of the fitness contribution of pollinator species $i$ to plant species $i$ (i.e., seeds produced per visit and per flower), in this case *Halimium calycinum*. Points indicate the posterior medians, while thick segments and thinner outer lines represent 50% and 90% intervals, respectively.

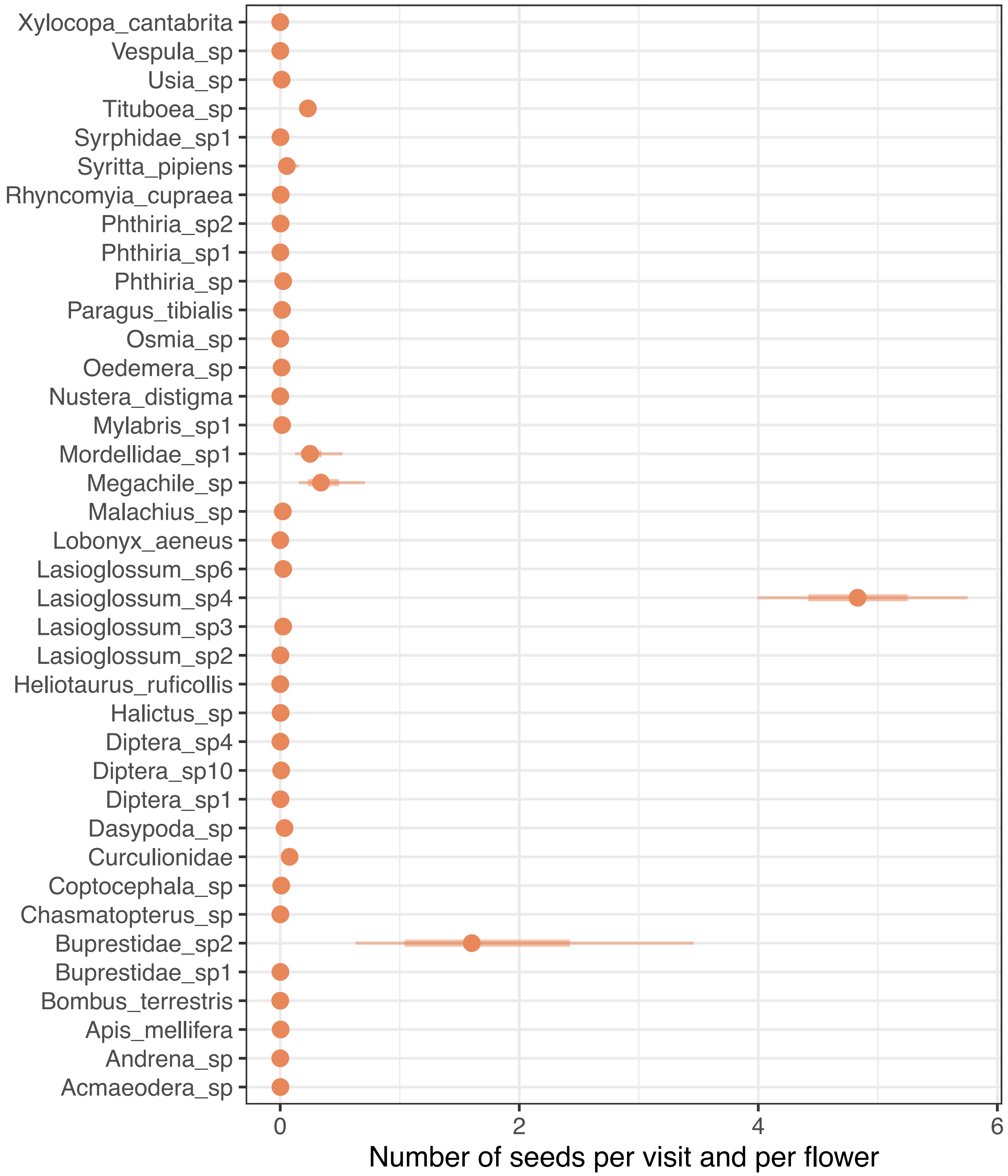


**Figure S3:** Credible intervals for parameters $\beta_{P_i A_i}$, which represent the value of the fitness contribution of pollinator species $i$ to plant species $i$ (i.e., seeds produced per visit and flower), in this case *Halimium halimifolium*. Points indicate the posterior medians, while thick segments and thinner outer lines represent 50% and 90% intervals, respectively.

### D Mean-field approximation to parameterize interaction coefficients

As described in the main text, we empirically parameterized $\gamma_{P_{is}A_i}$ in the interaction matrix $M$. To maintain focus only on the effects of the observed intraspecific plant variation in the mutualistic benefits received, all other coefficients in this interaction matrix were parameterized following a mean-field approximation. We set $\alpha_{P_{is}P_{it}}$ $(s = t) = \alpha_{A_iA_i} = 1$, $\alpha_{P_{is}P_{it}}$ $(s \neq t) = 0.9$, $\alpha_{P_{is}P_{jt}} = \alpha_{A_iA_j} = 0.1$, and $\gamma_{A_iP_{is}} = 0.2$ following previous work on the dynamics of mutualistic communities (Rohr *et al.*, 2014; Saavedra *et al.*, 2016; García-Callejas *et al.*, 2023). Because we combined empirically informed and mean-field parameterizations for different coefficients in the interaction matrix $M$, we do not actually know the realistic magnitude of intraspecific plant competition ($\alpha_{P_{is}P_{it}}$) relative to the magnitude of plant mutualistic benefits' ($\gamma_{P_{is}A_i}$). This is because the mean-field values of plant competition coefficients do not a priori have a biologically meaningful interpretation as it happens for the empirically estimated mutualistic benefits received by plant individuals. To account for the potential effects of the magnitude of plant competition, we explored the consequences of using different mean-field values for $\alpha_{P_{is}P_{it}}$ $(s \neq t)$ (Figs. S4, S5). We found that our qualitative results for the estimation of the size of the feasibility domain were robust to changes in the magnitude of intraspecific plant competition ($\alpha_{P_{is}P_{it}}$, $s \neq t$) while keeping empirical estimations for mutualistic benefits received by plant individuals ($\gamma_{P_{is}A_i}$) (Figs. S4, S5).

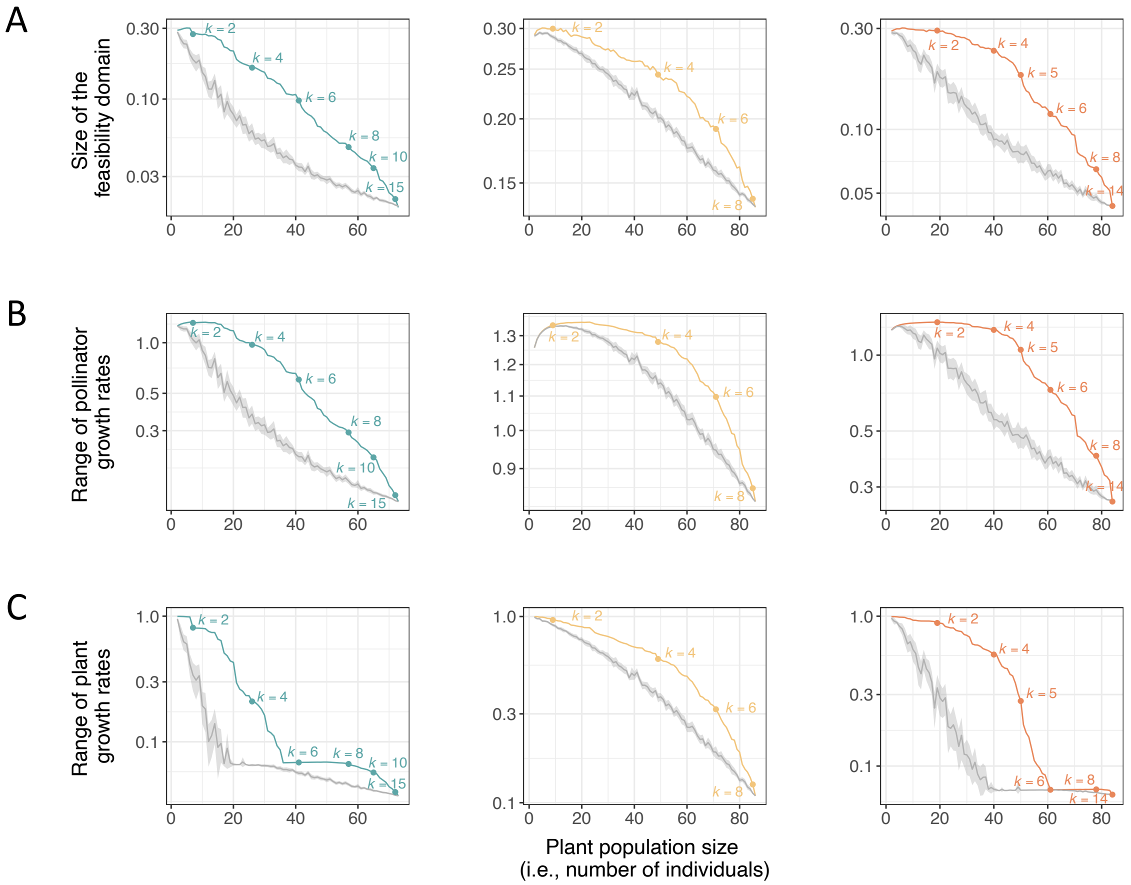


**Figure S4:** Effects of a higher proportion of specialized plant individuals in pollinator use, compared to mixtures of specialized and generalized plant individuals, on the persistence of plant populations' mutualistic assemblages, considering $\alpha_{P_{is}P_{it}}$ $(s \neq t) = 0.5$. For the three focal plant species (*Cistus libanotis* in blue; *Halimium calycinum* in yellow; and *Halimium halimifolium* in orange), gray lines represent the size of the feasibility domain of the entire assemblage (A), and the range of feasible growth rates of the plant species (B) and of the pollinator community (C) over an increasing population size by including plant individuals regardless of their degree. Colored lines represent the size of the feasibility domain of the entire assemblage (A), and the range of feasible growth rates of the plant species (B) and of the pollinator community (C) over an increasing plant population size by incorporating plant individuals from lower to a higher degree (i.e., from more specialized to more generalized individuals). Plant individuals with the same degree were included in order of decreasing specificity (i.e., decreasing coefficient of variation of interactions). Colored dots represent the degree ($k$) of plant individuals being included to increase population size. Differences in the size of feasibility domain and in the ranges of plant and pollinator feasible growth rates between the colored and gray lines for a given population size correspond to the effects of having a higher proportion of specialized individuals in the focal species, compared to a mixture of specialized and generalized individuals. Note that the lines corresponding to a higher proportion of specialized individuals (colored) are located above the confidence interval (95%) of the random mixtures of specialized and generalized (gray) across all population sizes.

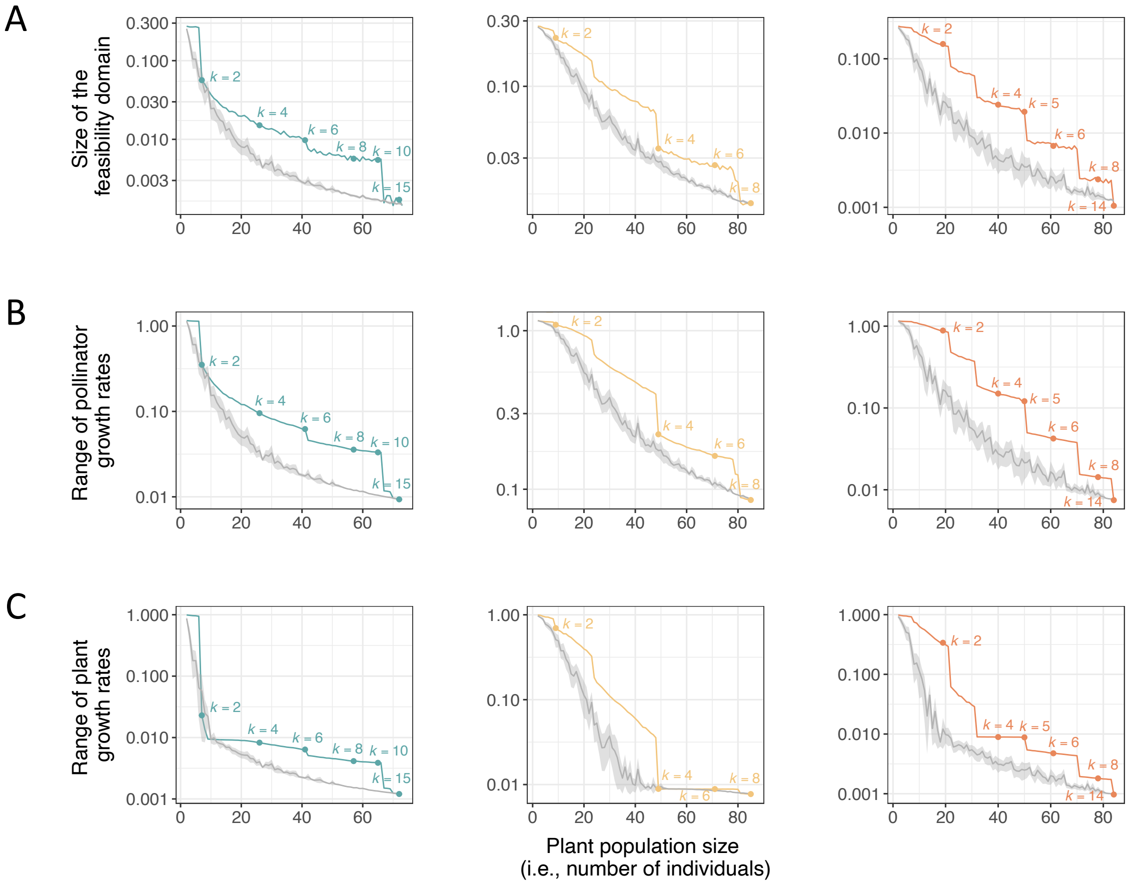


**Figure S5:** Effects of a higher proportion of specialized plant individuals in pollinator use, compared to mixtures of specialized and generalized plant individuals, on the persistence of plant populations' mutualistic assemblages, considering $\alpha_{P_{is}P_{it}}$ $(s \neq t) = 1.5$. For the three focal plant species (*Cistus libanotis* in blue; *Halimium calycinum* in yellow; and *Halimium halimifolium* in orange), gray lines represent the size of the feasibility domain of the entire assemblage (A), and the range of feasible growth rates of the plant species (B) and of the pollinator community (C) over an increasing population size by including plant individuals regardless of their degree. Colored lines represent the size of the feasibility domain of the entire assemblage (A), and the range of feasible growth rates of the plant species (B) and of the pollinator community (C) over an increasing plant population size by incorporating plant individuals from lower to a higher degree (i.e., from more specialized to more generalized individuals). Plant individuals with the same degree were included in order of decreasing specificity (i.e., decreasing coefficient of variation of interactions). Colored dots represent the degree ($k$) of plant individuals being included to increase population size. Differences in the size of feasibility domain and in the ranges of plant and pollinator feasible growth rates between the colored and gray lines for a given population size correspond to the effects of having a higher proportion of specialized individuals in the focal species, compared to a mixture of specialized and generalized individuals. Note that the lines corresponding to a higher proportion of specialized individuals (colored) are located above the confidence interval (95%) of the random mixtures of specialized and generalized (gray) across all population sizes.

### E Theoretical testing of the effects of intraspecific variation

To assess how intraspecific plant variation in interaction patterns with pollinators impact feasibility estimates in a simplified community, we used different abundances for pollinator species ($A_1$=0.4, $A_2$=0.6) and parameterized interaction coefficients as follows. We set $\alpha_{P_{11}P_{11}} = \alpha_{P_{12}P_{12}} = 1$, $\alpha_{P_{11}P_{12}} = \alpha_{P_{12}P_{11}} = 0.9$, $\alpha_{A_1A_1} = \alpha_{A_2A_2} = 1$, $\alpha_{A_1A_2} = \alpha_{A_2A_1} = 0.1$, and $\gamma_{A_1P_{11}} = \gamma_{A_1P_{12}} = \gamma_{A_2P_{11}}$ $= \gamma_{A_2P_{12}} = 0.2$. The mutualistic benefits received by plant individual types from pollinator species ($\gamma_{P_{11}A_1}$, $\gamma_{P_{11}A_2}$, $\gamma_{P_{12}A_1}$, $\gamma_{P_{12}A_2}$) were parameterized following Eq. 8 in Box 1, considering $\beta_{P_1A_1}$=0.4, $\beta_{P_1A_2}$=0.2, spanning different values of visitation rates ($v_{P_{11}A_2}$, $v_{P_{11}A_2}$, $v_{P_{12}A_2}$ and $v_{P_{12}A_2}$) (Fig. 1C), and assuming equal flower production across plant individual types ($f_{P_{11}}$= $f_{P_{12}}$).

To isolate the effects of differences in pollinators' contributions from those of differences in their abundances, we tested under which conditions a system can be feasible when pollinator species are equally abundant ($A_1$=$A_2$=0.5). In this case, when pollinators contribute equally to plant fitness, the highest feasibility is achieved when the proportion of visits received by plant types are balanced between both pollinator species (i.e., the number of visits by both pollinator species are complementary) (Fig. S7A). In this case, the system can be feasible even if both plant types are highly specialized on a different pollinator species, which is the maximum intraspecific variation possible. In contrast, as the difference in fitness contribution between the pollinator species increases, a strong specialization of any given plant individual type to a given pollinator species drastically reduces the feasibility of this system (Fig. S7B-C), as the plant type specialized on the more effective pollinator species (i.e., that with the highest contribution to plant fitness) will outcompete the other plant type. These results underscore the consequences of the presence of pollinator species differing in their contributions to plant fitness, as observed in real-world systems, as these differences impose strong constraints on how specialized plant individual types can be to avoid intraspecific competitive exclusion.

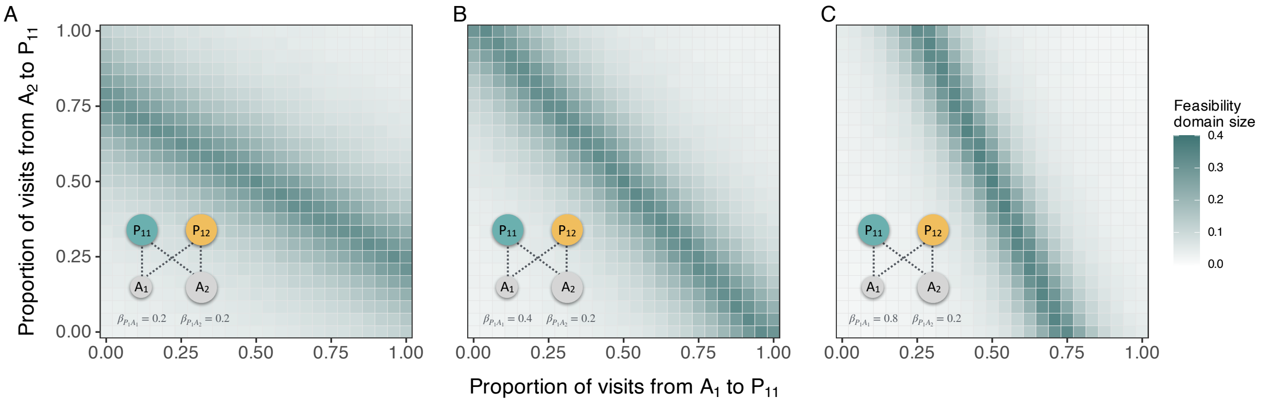


**Figure S6:** Variation in the feasibility domain size depending on the proportion of visits from two pollinator species ($A_1$ and $A_2$) to plant individual type $P_{11}$ (dotted links in the graph) when pollinator species differ in abundance ($A_1 < A_2$). We assumed that (A) both pollinator species contribute equally to plant fitness ($\beta_{P_1}{=}\beta_{P_2}$), (B) one pollinator species contributes twice as much as the other to plant fitness, and (C) one pollinator species contributes four times as much as the other to plant fitness. The 'fully specialized' case is represented in the top left corner, where $P_{11}$ is specialized on $A_2$ and $P_{12}$ on $A_1$, or the bottom right corner, where $P_{11}$ is specialized on $A_1$ and $P_{12}$ on $A_2$. The 'fully generalized' case is represented in the center of the grid, where both plant types receive the same exact proportion of visits. All other cells in the grid represent intermediate cases between mixtures of generalized and specialized types.

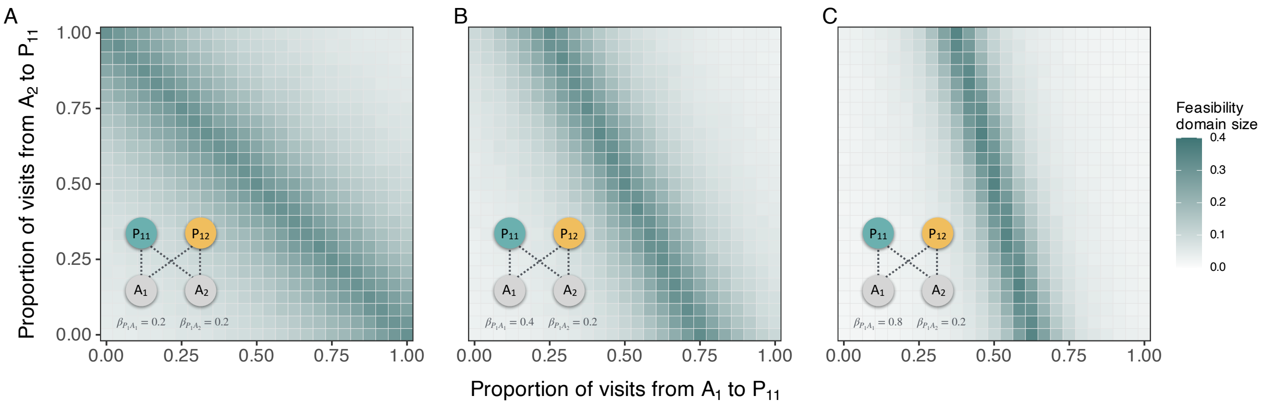


**Figure S7:** Variation in the feasibility domain size depending on the proportion of visits from two pollinator species ($A_1$ and $A_2$) to plant individual type $P_{11}$ (dotted links in the graph) when pollinator species have the same abundance ($A_1 = A_2$). We assumed that (A) both pollinator species contribute equally to plant fitness ($\beta_{P_1}$=$\beta_{P_2}$), (B) one pollinator species contributes twice as much as the other to plant fitness, and (C) one pollinator species contributes four times as much as the other to plant fitness. The 'fully specialized' case is represented in the top left corner, where $P_{11}$ is specialized on $A_2$ and $P_{12}$ on $A_1$, or the bottom right corner, where $P_{11}$ is specialized on $A_1$ and $P_{12}$ on $A_2$. The 'fully generalized' case is represented in the center of the grid, where both plant types receive the same exact proportion of visits. All other cells in the grid represent intermediate cases between mixtures of generalized and specialized types.

### F Components influencing plant individuals' fitness

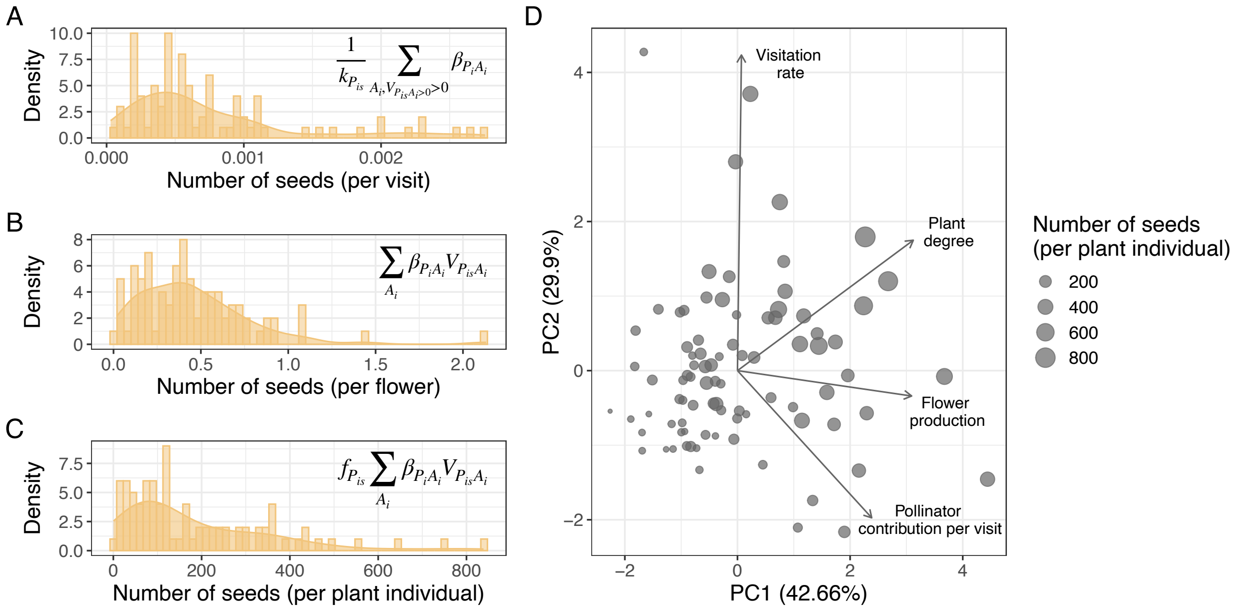


**Figure S8:** Intraspecific plant fitness variation within one of the study species (*Halimium calycinum*) when incorporating different components influencing plant fitness. (A) Histogram of the number of seeds produced per visit and flower across plant individuals, where intraspecific variation is promoted by differences in pollinator assemblages and their associated contribution to fitness. (B) Histogram of the number of seeds produced per flower across plant individuals, where intraspecific variation is promoted by differences in visitation rates in addition to differences in pollinator assemblages. (C) Histogram of the total number of seeds contributed to the population seed bank across plant individuals, where intraspecific variation is produced by differences in pollinator assemblages, visitation rates and flower production. (D) Principal Component Analysis (PCA) of plant individuals (dots) based on different sources of intraspecific variation influencing plant fitness. Dot size is proportional to the number of seeds produced and contributed to the seed bank of the population. Estimates of the number of seeds per visit (A), per flower (B) and per individual plant (C) were calculated following the equations shown in each panel, in which $k_{P_{is}}$ denotes de degree of plant individual $P_{is}$, $\beta_{P_i A_i}$ denotes the number of seeds per flower produced in plant species $P_i$ per visit of pollinator species $A_i$, $v_{P_{is} A_i}$ is the visitation rate per flower of pollinator species $A_i$ to plant individual $P_{is}$, and $f_{P_{is}}$ represent the number of flowers produced by plant individual $P_{is}$.

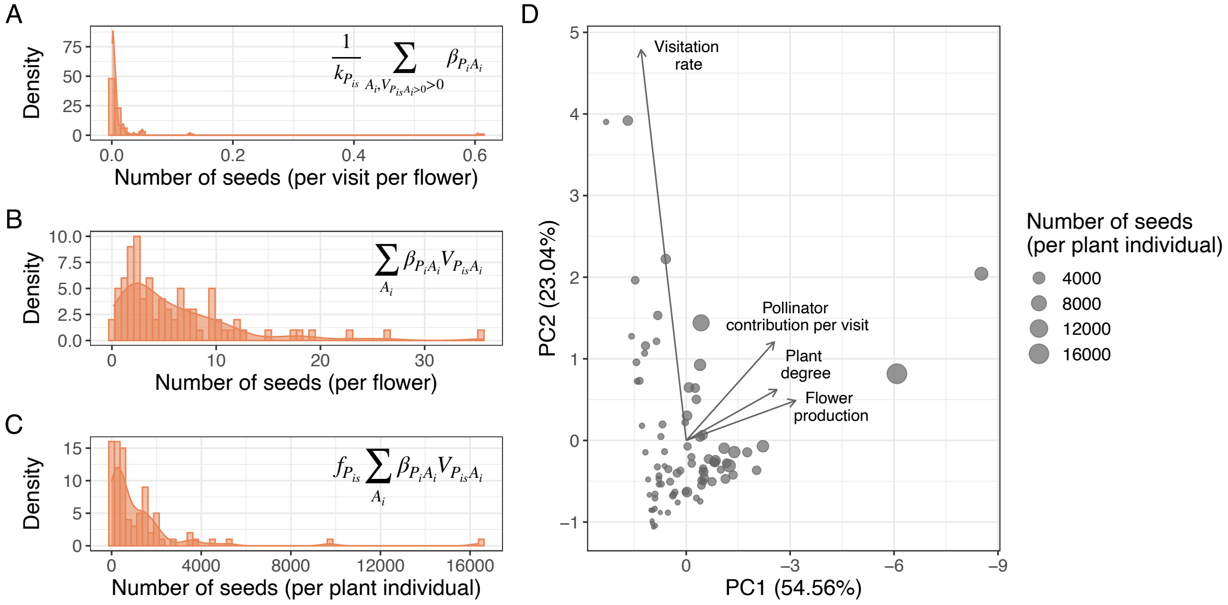


**Figure S9:** Intraspecific plant fitness variation within one of the study species (*Halimium halimifolium*) when incorporating different components influencing plant fitness. (A) Histogram of the number of seeds produced per visit and flower across plant individuals, where intraspecific variation is promoted by differences in pollinator assemblages and their associated contribution to fitness. (B) Histogram of the number of seeds produced per flower across plant individuals, where intraspecific variation is promoted by differences in visitation rates in addition to differences in pollinator assemblages. (C) Histogram of the total number of seeds contributed to the population seed bank across plant individuals, where intraspecific variation is produced by differences in pollinator assemblages, visitation rates and flower production. (D) Principal Component Analysis (PCA) of plant individuals (dots) based on different sources of intraspecific variation influencing plant fitness. Dot size is proportional to the number of seeds produced and contributed to the seed bank of the population. Estimates of the number of seeds per visit (A), per flower (B) and per individual plant (C) were calculated following the equations shown in each panel, in which $k_{P_{is}}$ denotes de degree of plant individual $P_{is}$, $\beta_{P_i A_i}$ denotes the number of seeds per flower produced in plant species $P_i$ per visit of pollinator species $A_i$, $v_{P_{is} A_i}$ is the visitation rate per flower of pollinator species $A_i$ to plant individual $P_{is}$, and $f_{P_{is}}$ represent the number of flowers produced by plant individual $P_{is}$.

### G Relationship between feasibility and fitness components of plant individual types

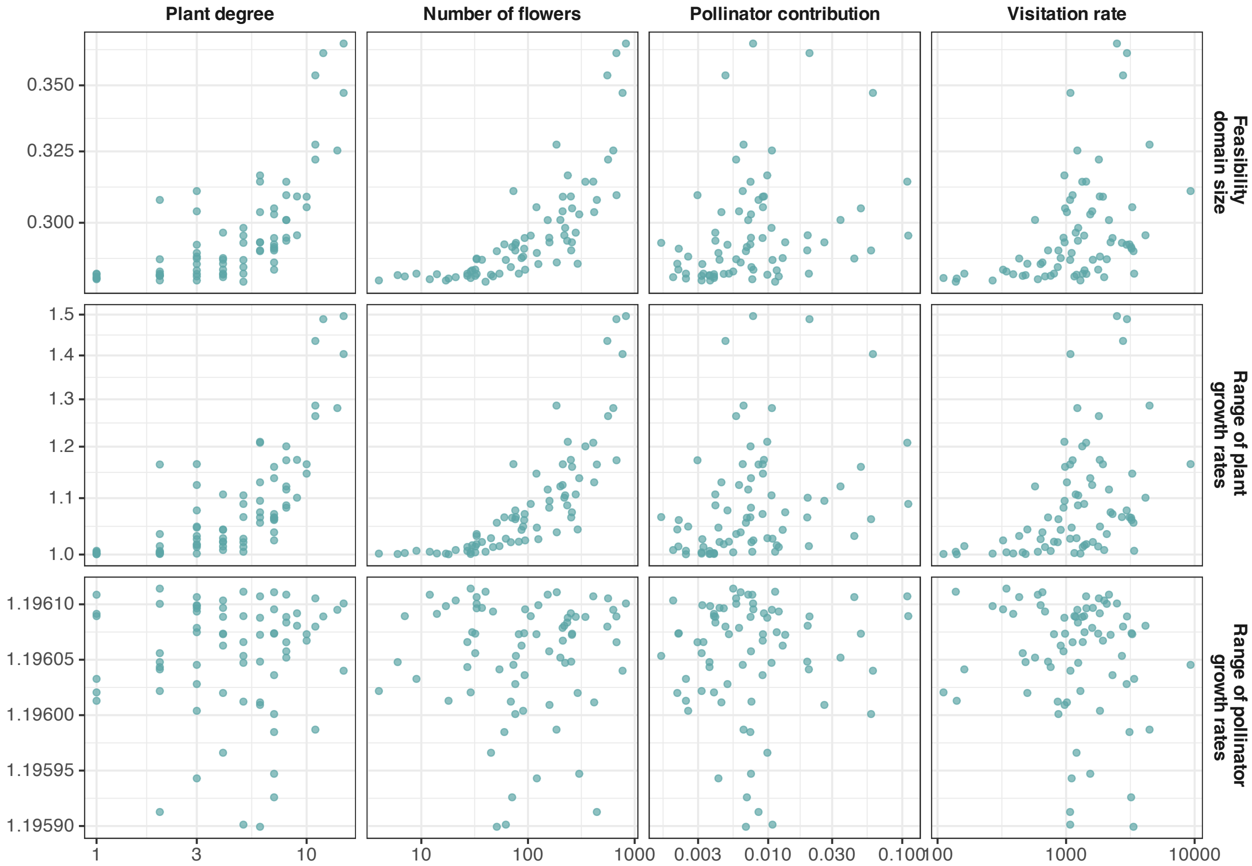


**Figure S10:** Impact of different components influencing the fitness of plant individual types (i.e., represented by sampled plant individuals in the empirical case study) on the feasibility domain size of a mutualistic assemblage composed exclusively of that plant type within a *Cistus libanotis* population, assuming no intraspecific variation. Each dot represents the plant individual type composing each mutualistic assemblage. Therefore, dots illustrate how different attributes of a given plant type impact feasibility when the plant population is composed only by individuals belonging to that type.

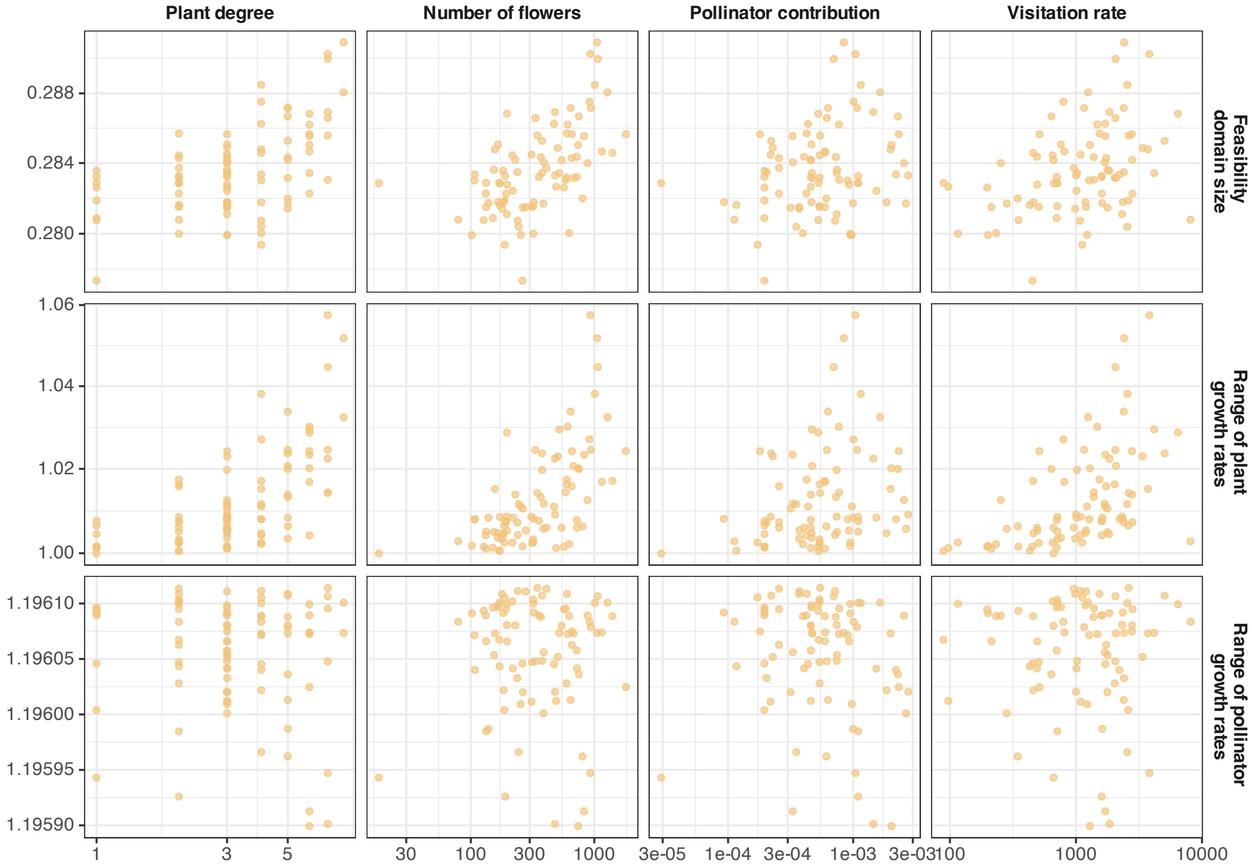


**Figure S11:** Impact of different components influencing the fitness of plant individual types (i.e., represented by sampled plant individuals in the empirical case study) on the feasibility domain size of a mutualistic assemblage composed exclusively of that plant type within a *Halimium calycinum* population, assuming no intraspecific variation. Each dot represents the plant individual type composing each mutualistic assemblage. Therefore, dots illustrate how different attributes of a given plant type impact feasibility when the plant population is composed only by individuals belonging to that type.

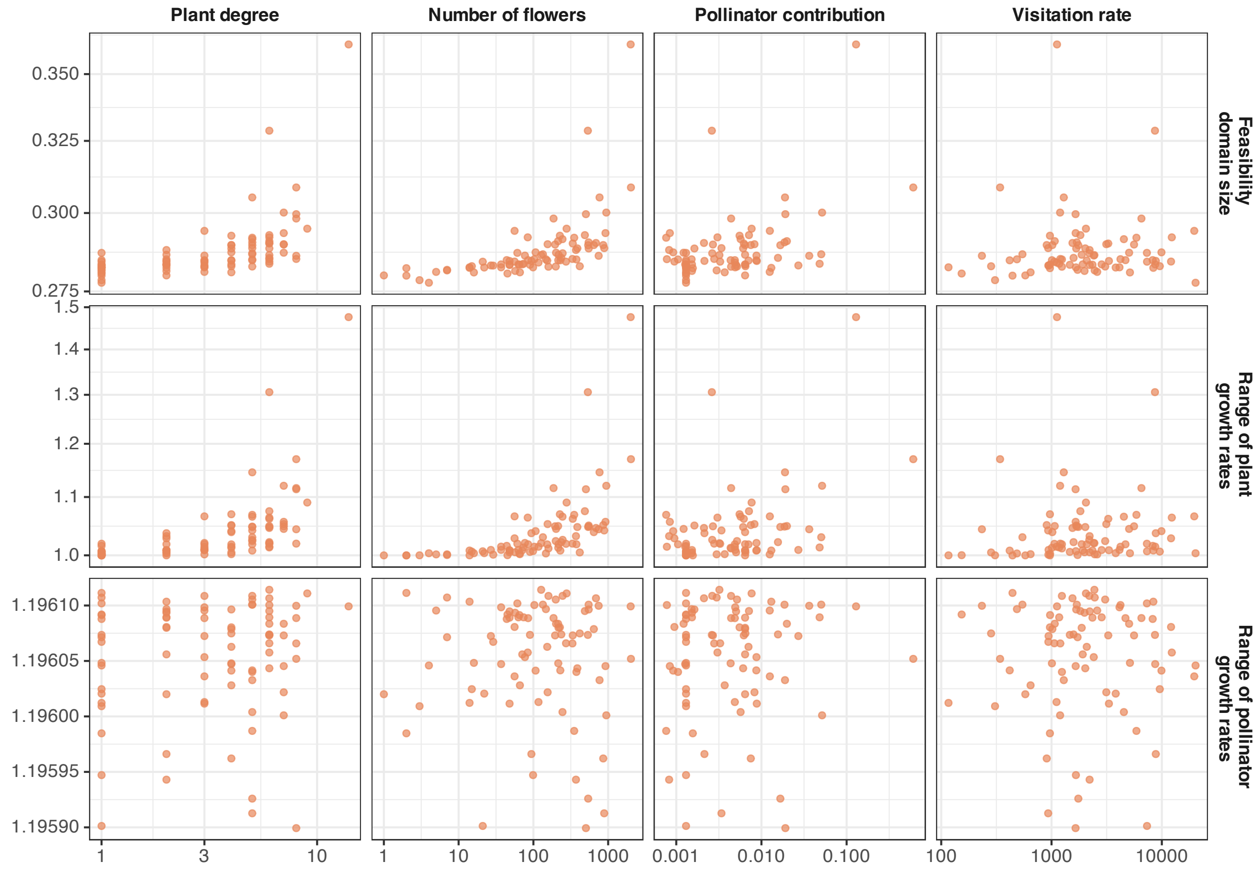


**Figure S12:** Impact of different components influencing the fitness of plant individual types (i.e., represented by sampled plant individuals in the empirical case study) on the feasibility domain size of a mutualistic assemblage composed exclusively of that plant type within a *Halimium halimifolium* population, assuming no intraspecific variation. Each dot represents the plant individual type composing each mutualistic assemblage. Therefore, dots illustrate how different attributes of a given plant type impact feasibility when the plant population is composed only by individuals belonging to that type.
